# Real-time monitoring of mechanical cues in the regenerative niche reveal dynamic strain magnitudes that enhance bone repair

**DOI:** 10.1101/663278

**Authors:** Brett S. Klosterhoff, Jarred Kaiser, Bradley D. Nelson, Salil S. Karipott, Marissa A. Ruehle, Scott J. Hollister, Jeffrey A. Weiss, Keat Ghee Ong, Nick J. Willett, Robert E. Guldberg

**Affiliations:** George W. Woodruff School of Mechanical Engineering, Georgia Institute of Technology, Atlanta, GA; Parker H. Petit Institute for Bioengineering and Bioscience, Georgia Institute of Technology, Atlanta, GA; Research Service, Atlanta VA Medical Center, Decatur, GA; Department of Orthopaedics, Emory University, Atlanta, GA; Department of Biomedical Engineering, Michigan Technological University, Houghton, MI; Wallace H. Coulter Department of Biomedical Engineering, Georgia Institute of Technology and Emory University, Atlanta, GA; Department of Biomedical Engineering, University of Utah, Salt Lake City, UT; Scientific Computing and Imaging Institute, University of Utah, Salt Lake City, UT; Department of Orthopedics, University of Utah, Salt Lake City, UT; Knight Campus, University of Oregon, Eugene, OR

## Abstract

Mechanical loads exerted on the skeleton during activities such as walking are important regulators of bone repair, but dynamic biomechanical signals are difficult to measure inside the body. The inability to measure the mechanical environment in injured tissues is a significant barrier to developing integrative regenerative and rehabilitative strategies that can accelerate recovery from fracture, segmental bone loss, and spinal fusion. Here we engineered an implantable strain sensor platform and measured strain across a bone defect in real-time throughout rehabilitation. We used the sensor to longitudinally quantify mechanical cues imparted by a load-sharing fixation plate that significantly enhanced bone regeneration in rats. We found that sensor readings correlated with the status of healing, suggesting a potential role for strain sensing as an X-ray-free healing assessment platform. This study demonstrates a promising approach to quantitatively develop and exploit mechanical rehabilitation strategies that enhance bone repair.

## Introduction

Tissue regeneration requires dynamic and spatially coordinated cellular activity involving bidirectional interactions between the healing niche and surrounding environment. Characterizing the progression of early-stage environmental cues in the regenerative niche and their contribution to divergent healing outcomes is a critical and active area of research. An improved understanding of biochemical and biophysical cues may inform the development of therapies that modulate the microenvironment to more effectively resolve challenging injuries (*1*). Mechanical signals are among the most potent and dynamic environmental regulators of tissue repair (*2*, *3*). In particular, forces exerted on musculoskeletal tissues during locomotion strongly influence regenerative processes (*4*, *5*). The sensitivity of bone to extrinsic mechanical cues is well documented, where a moderate magnitude of mechanical loading accelerates osteoprogenitor differentiation, matrix mineralization, and restoration of biomechanical function (*6*–*8*).

Strategies for controlled transfer of mechanical loads in vivo are a promising therapeutic target to promote osteogenesis after fracture, segmental bone loss, spinal fusion, and joint arthroplasty. Indeed, there is substantial clinical need to enhance bone regeneration for the millions of patients undergoing these procedures each year, as a significant sub-set are afflicted with prolonged disability or multiple revision surgeries due to non-union or poor osseointegration (*9*). Mechanical stimulation strategies have spanned a range of approaches and garnered significant scientific and clinical interest including external or percutaneous fixators and actuators (*10*), and stabilization hardware or scaffolds with reduced stiffness to permit load sharing (*11*–*13*). Additionally, more functionally relevant, non-invasive methods such as physical rehabilitation and exercise may be implemented to enhance load transfer while simultaneously inhibiting adjacent tissue atrophy and accelerating functional recovery of daily activities (*14*–*16*). Conversely, excessive loading can impair healing resulting in fibrosis, hypertrophic non-union, or construct failure (*8*, *17*). A critical challenge limiting these techniques has been the ability to monitor the dynamic mechanical environment at the regenerative niche during repair—this mechanical environment has rarely been quantified, resulting in incomplete understanding of biomechanical conditions and limited means with which to reliably investigate the safety and efficacy of mechanical based interventions.

A promising approach to quantitatively evaluate mechanical cues during tissue repair is the integration of electromechanical sensors with devices already typically installed at the site of healing, such as fixation plates, implants, or scaffolds (*18*). Advancements in microfabrication and wireless data transfer have attained a level of maturity and a sufficiently small size for biomedical applications. Recent reports of a broad range of transient and permanent implantable sensors have facilitated new physiological insights in pre-clinical in vivo models as well as digital health monitoring in humans (*19*–*21*).

Using a pre-clinical rat model of long bone repair, we observed that varying load transfer modulates vascular and skeletal tissue formation after injury (*7*, *17*). Motivated to better understand the temporal progression of tissue-level mechanical cues that could enhance skeletal repair, we recently developed a fully implantable strain sensor platform enabling real-time monitoring of mechanical boundary conditions across the defect during functional rehabilitative activities like walking (*22*). The device possesses sufficient sensitivity and size for implementation in rats--animals which are used commonly used as a key pre-clinical test bed to screen therapeutic approaches before scaling up to more costly large animal models.

Here, we deployed this strain sensor platform with the objective to examine the evolution of biomechanical cues in the regenerative niche after injury and assess the effects of differing magnitudes of functional mechanical loading on bone defect repair and revascularization. We hypothesized that a moderate increase in ambulatory load sharing conferred by reduced stiffness fixation would increase mechanical stimuli initially, and that the elevated deformation would eventually decrease due to enhanced bone formation. We found that rehabilitative load sharing permitted by reduced stiffness of fixation substantially accelerated and enhanced bone repair. Furthermore, we observed that real-time monitoring of strain magnitude during gait correlated with the status of bone repair and that osteogenic ambulatory loading differentially regulated neovascular growth after injury. These results demonstrate the potential of advancements in biomedical sensors to optimize mechanobiological therapies by remotely monitoring dynamic biophysical cues in vivo.

## Results

### Real-time and on-demand ambulatory monitoring of in vivo mechanical boundary conditions across bone defects using an integrated strain sensor

To assess the relative effects of mechanical loading on bone repair, we used an established rat femoral bone defect model with well-characterized healing kinetics (*23*). The critically-sized 6 mm defects received one of two treatments: 2 µg bone morphogenetic protein-2 (BMP-2), a minimal osteogenic dose that typically results in defects at the threshold of mineralized bridging at 8 weeks, or left empty as non-healing negative controls. The mechanical environment under ambulatory loads was perturbed by stabilizing defects with either stiff or moderately compliant internal fixation plates (fixators) which possessed a modular bridging segment fabricated from polysulfone (PSU) or ultra-high molecular weight polyethylene (UHMWPE), respectively. Our previous work in this model demonstrated that early ambulatory loading in a highly axially compliant fixator which was 80% more compliant than stiff PSU fixators drastically inhibited bone and vascular repair due to excessive deformation (*17*). In this study, we sought to explore a more moderate mechanical environment that could enhance bone regeneration. Therefore, compliant UHMWPE fixators featured a flexural stiffness about 40% lower than stiff PSU counterparts of identical geometry (Fig. 1A; UHMWPE = 145.2 ± 10.6, PSU= 232.4 ± 20.1). Full study timelines are outlined in Table S1.

**Fig. 1:**
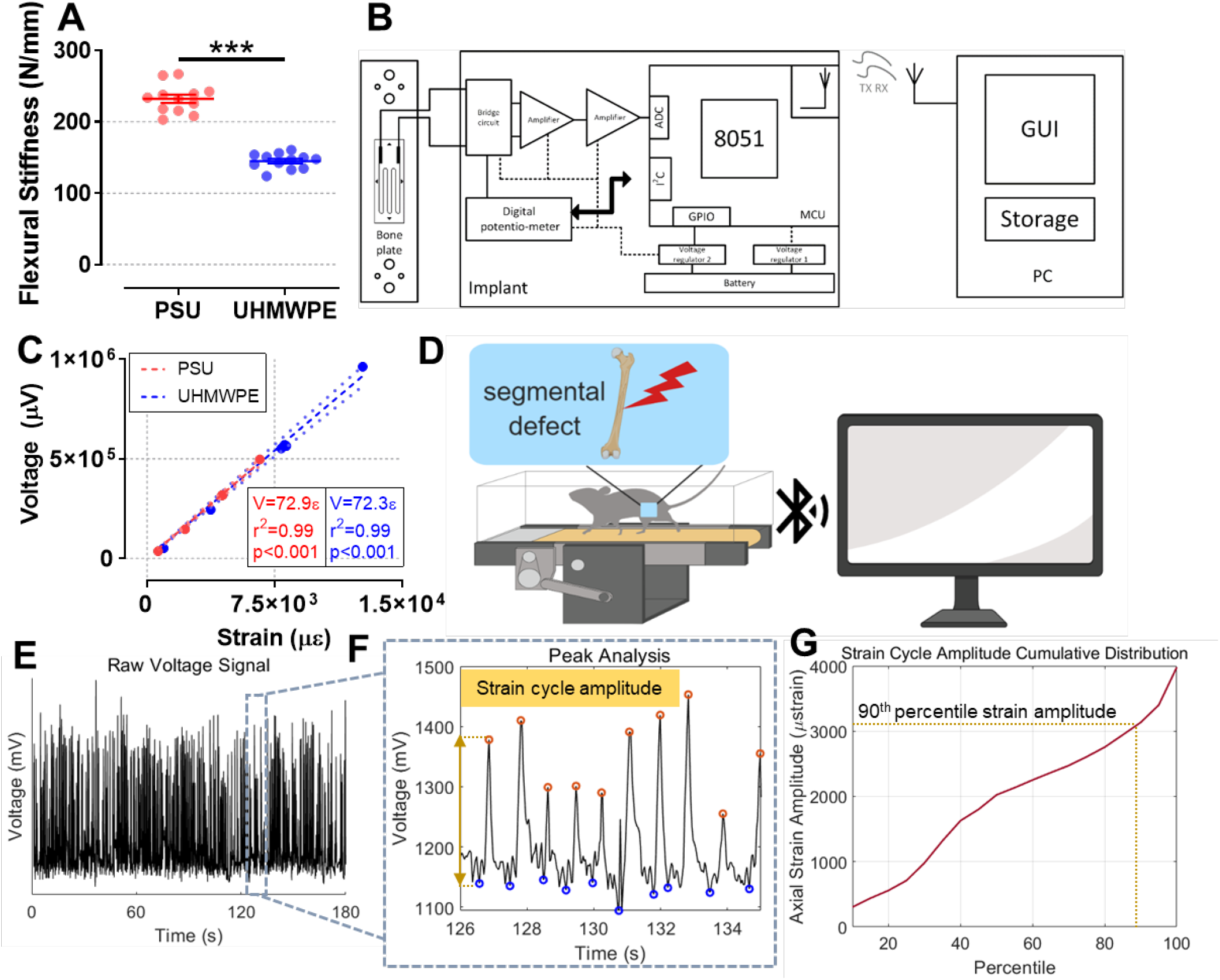
Real-time and on-demand ambulatory monitoring of in vivo mechanical boundary conditions across segmental bone defects using an integrated strain sensor. **(A)** UHMWPE fixation plates possessed a flexural stiffness 40% lower than PSU. n = 12. ***p < 0.001 via t-test. **(B)** Functional block diagram depicting fixator-mounted strain sensor interfaced with intra-abdominal transceiver unit consisting of analog front-end, MCU, GPIO, and BLE interface providing transmit and receive functionality with a host computer. **(C)** Representative electromechanical calibrations for UHMWPE and PSU fixators, strain sensors exhibited high linearity and precision. n = 20 cycles per strain. r^2^ > 0.99, ***p < 0.001 via Pearson’s. **(D)** Rats walked on a treadmill twice weekly 7 days after surgical creation of a unilateral segmental femoral defect and strain measurements were acquired in real-time (See Movie S1). **(E)** Experimental sensor output during a 3 minute recording of rodent gait **(F)** Inset of sensor output depicting individual strain cycles corresponding to discrete steps and corresponding amplitudes. **(G)** Experimental cumulative distribution of strain cycles recorded during a treadmill session, indicating the 90^th^ percentile amplitude.

In vivo strain measurements were facilitated via the integration of a strain sensor into the polymeric bridging segments of each fixator. The sensor transmitted real-time uniaxial strain measurements to a remote laptop with a USB receiver up to five meters away via a Bluetooth Low Energy (BLE) enabled digital transceiver (Fig. 1B). The transceiver was remotely controlled to either transmit data or enter a low-power receiving mode by commands from the computer, enabling on-demand user control of data collection and power allocation. All devices exhibited linear responses and sufficient sensitivity to detect applied strain throughout the physiological dynamic range (Fig. 1C).

Ambulatory loading is a promising and straightforward approach to administer dynamic biophysical stimuli to the regenerative niche and enhance tissue repair (*16*). Therefore, we sought to assess the temporal progression of mechanical cues imparted during slow walking by measuring strain during treadmill sessions starting one week after surgery and continuing twice weekly thereafter. Strain acquisition proceeded biweekly through 5 weeks and once weekly thereafter (Fig. 1D). Dynamic strain cycle amplitudes corresponding to individual steps were computed and ranked by magnitude (Fig. 1E-G). The 90^th^ percentile strain magnitude was tracked longitudinally as it represented a threshold of the 50-60 highest magnitude strain cycles, a sufficient number of cycles to provoke an adaptive cellular response (*6*, *24*, *25*). Rehab collection periods represented the most significant mechanical stimulus exerted on the femur in terms of both magnitude and frequency content; treadmill walking produced a significant 60% increase in strain magnitude relative to nocturnal in-cage activity (Fig. S1). To demonstrate real-time strain measurements during ambulatory activity with high temporal resolution, high-speed x-ray video of an animal walking on a treadmill during simultaneous strain acquisition was recorded one month after surgery (Movie S1 & Fig. S2). Video analysis demonstrated variations in strain signal were synchronized with stance and swing phases of walking, where stance phases produced a transient flexural strain on the fixator, conferring a compressive environment on the healing tissue. These data demonstrate the utility of implantable sensor technologies to remotely quantify mechanobiological cues within regenerating in vivo environments at meaningfully protracted time points after implantation.

### Rehabilitative load sharing increased mechanical stimulation and accelerated bridging of segmental bone defects

Sensor measurements of longitudinal in vivo strains confirmed that compliant UHMWPE fixators permitted higher magnitude mechanical stimulation in vivo. Initial strain magnitudes were increased by approximately two-fold compared to stiff PSU fixators (Fig. 2A). Strain magnitudes did not change appreciably throughout the 8 week study in empty defect non-healing controls, regardless of fixator stiffness. Strain magnitudes on defects treated with a low dose of BMP-2 and stabilized by compliant UHMWPE fixators diverged from corresponding UHMWPE empty controls beginning at 2 weeks and continued to decline gradually until converging with levels observed in stiff PSU fixators. A similar decline was not observed in BMP-2 treated defects stabilized by PSU fixators, which showed no significant changes throughout the duration of the study. Serial radiographs corroborated our hypothesis that the progressive strain decline observed in UHMWPE fixators was due to bridging of mineralized tissue across the segmental defect, consequently reducing load share carried by the fixator (Fig. 2B & C). Between weeks 2 and 4, the average strain magnitude in the UHMWPE group was halved from 5408 ± 704 µε to 2698 ± 567 µε, coinciding with a marked 64% increase in the percentage of bridged defects.

**Fig. 2:**
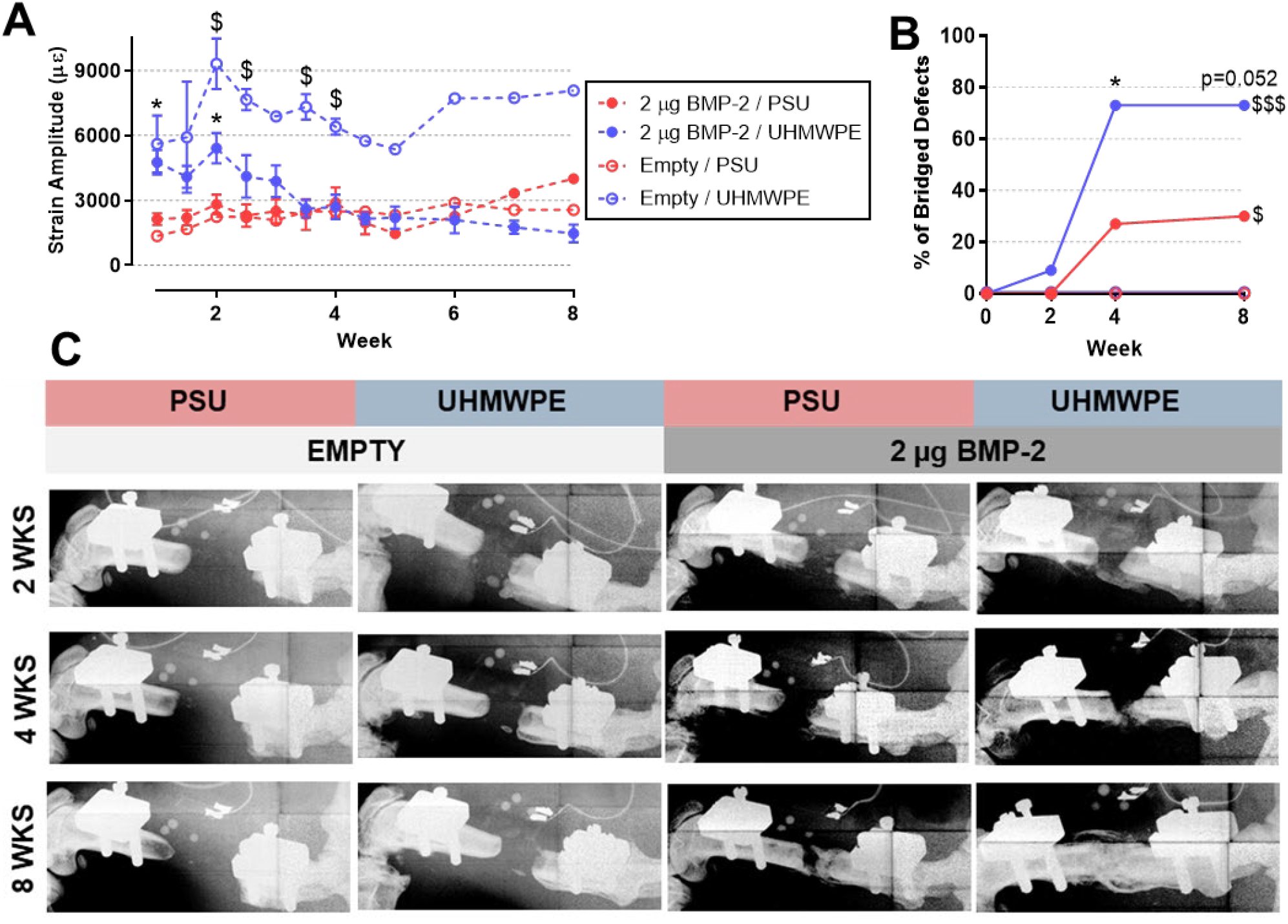
Rehabilitative load sharing initially increased mechanical stimulation and accelerated bridging of segmental bone defects. **(A)** Longitudinal strain amplitude measurements verified UHMWPE fixation permit increased mechanical stimulation of bone defects. n=1-9. 2 µg BMP-2: *p < 0.05 UHMWPE vs. PSU. UHMWPE: $p < 0.05 Empty vs 2 µg BMP-2 via Two-way ANOVA with Tukey’s test. **(B)** Longitudinal analysis of bone bridging via in vivo microCT segmented at 50% intact cortical bone mineral density. n = 10-11. *p < 0.05 differences between groups via chi-square test. $p < 0.05 significance of trend via chi-square test for trend. **(C)** Representative longitudinal x-ray images for each group, demonstrating substantially increased mineralization with compliant UHMWPE fixators in the presence of BMP-2.

A fundamental motivation for developing the strain sensor platform was to elucidate dynamic mechanical boundary conditions in a healing skeletal defect and to identify specific ranges that could enhance repair. Radiographic comparisons between groups demonstrated that rehabilitative load sharing permitted by UHMWPE fixators significantly accelerated bridging and enhanced total mineralization in the presence of BMP-2. Bridging ratios were nearly tripled in UHMWPE stabilized defects compared to PSU at 4 weeks. Empty non-healing controls confirmed that negligible mineralized tissue developed without intervention in this critically-sized defect model irrespective of fixation stiffness, which correlates with a similar lack of temporal variation in strain magnitude. Facilitated by the implantable strain sensor, these data support the potential for controlled biophysical signals imparted by normal movement activities to accelerate tissue regeneration in challenging injuries.

### Bone regeneration was enhanced by ambulatory mechanical loading and initial strain magnitudes correlated with improved healing outcomes

We performed serial in vivo microcomputed tomography (microCT) scans to quantitatively assess the effect of the mechanical environment on bone formation. In agreement with the radiographic data, we observed a beneficial effect of mechanical loading on bone regeneration (Fig. 3A). Compliant UHMWPE fixation had significant overall effects on bone volume, trabecular thickness, trabecular number, and trabecular separation (Fig. 3B, D-F); at the 8 week time point bone volume and trabecular thickness in the UHMWPE group were significantly increased by 63% and 47%, respectively, compared to stiff PSU fixators. Mineral density and polar moment of inertia were not affected by load sharing (Fig. 3C & Fig. S3), suggesting that the early stage effects of mechanical stimulation are mediated primarily by accumulation of relatively centralized woven bone volume rather than acceleration of bone remodeling and mineral densification, processes whose kinetics may act over longer time spans. Picro Sirius Red histological analysis of 8 week tissue samples supported that UHMWPE fixation supported a mixture of woven and lamellar extracellular matrix (ECM), whereas PSU fixation exhibited primarily lamellar ECM (Fig. S4). Ex vivo torsion testing to failure yielded no significant effects on failure strength, though a single high-performing statistical outlier stabilized by PSU may have obscured the results, albeit this sample had no experimental observations to warrant exclusion (Fig. 3G).

**Fig. 3:**
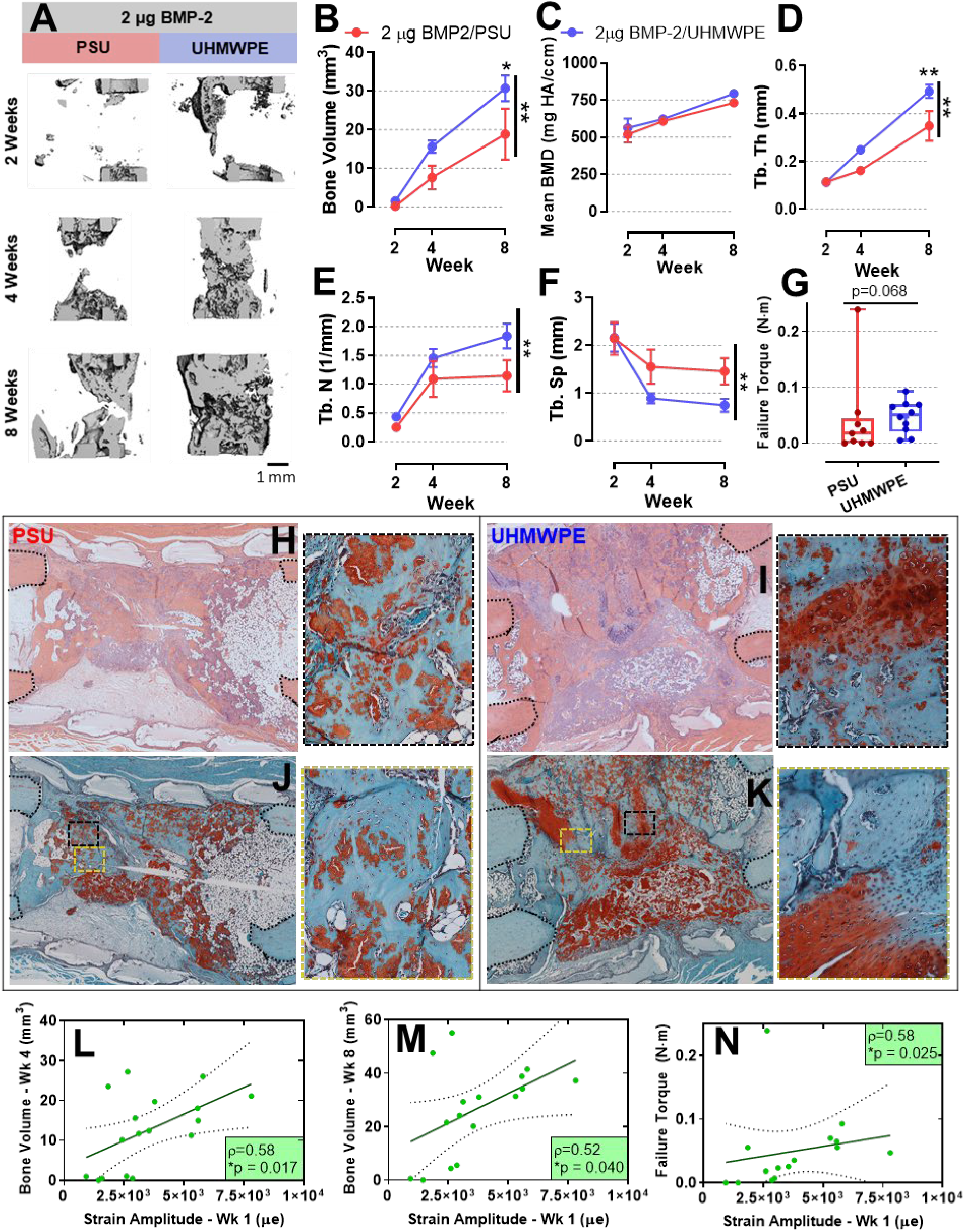
Bone regeneration was enhanced by ambulatory mechanical loading and initial strain amplitudes correlated positively with improved healing outcomes. **(A)** Bone defect microCT reconstructions representing median samples, showing increased mineralization with compliant UHMWPE fixation. Scale bar, 1 mm. Longitudinal microCT quantifications of **(B)** bone volume, **(C)** mean bone mineral density, **(D)** trabecular thickness, **(E)** trabecular number, **(F)** trabecular spacing demonstrating increased architectural parameters and no effects on mineral density with compliant UHMWPE fixators. n = 10-11. Vertically oriented bar ** p < 0.01 overall main effect UHMWPE vs. PSU via Two-way ANOVA. Overhead asterisks * p < 0.05, ** p < 0.01 via Sidak’s multiple comparisons test. **(G)** Failure torque of explanted femoral defects 8 weeks after surgery. n = 10-11. p = 0.068 via Mann-Whitney U test. **(H-I)** Week 8 H&E-stained histological sections showing intact femoral ends (black dotted lines) and increased mineralized tissue formation under compliant UHMWPE fixation. **(J-K)** Week 8 Safranin-O/Fast green sections demonstrate extensive regions of hypertrophic chondrocytes and endochondral bone formation under compliant UHMWPE fixation. Scale bars, 500 µm for full defect and 50 µm for insets. Specimen-specific strain amplitudes during the first treadmill activity period at 1 week, before appreciable mineralization had occurred, exhibited significant positive correlations with **(L)** 4 week bone volume, **(M)** 8 week bone volume **(N)** and 8 week failure torque. n = 17. *p < 0.05 via rank-order correlation.

Histological analysis of the 8 week samples revealed direct contact between the regenerating tissue and biomaterial irrespective of fixation stiffnesses, with regions of mineralized tissue surrounding remnants of alginate hydrogel (Fig. 3H-I). Safranin-O/Fast green staining indicated that compliant UHMWPE fixation exhibited extensive regions of hypertrophic chondrocytes and endochondral ossification throughout the bone defect, while chondrocytes were observed to a qualitatively lesser extent with PSU (Fig. 3J-K). These data are supported by prior reports that moderate mechanical loading primarily mediates osteogenesis via endochondral bone formation, the primary mechanism for long bone formation and fracture repair under load-bearing conditions (*5*, *6*, *8*, *17*, *26*).

The significant enhancement of bone regeneration stabilized by UHMWPE fixation supported further inquiry into the therapeutic potential of early, moderate mechanical stimulation. To evaluate this effect more carefully, we examined the relation between initial strain amplitudes and longer-term healing outcomes on a per animal basis to investigate if slight inter-animal variations in strain correlated with differential healing outcomes at disparate time points. Rank-order correlation revealed a significant positive relationship between strain cycle amplitude at week 1 and bone volume at weeks 4 and 8 and failure torque (Fig. 3L-N). This correlation with tissue strain was no longer significant at 2 weeks (Fig. S5A-C); radiographic shadowing and mineralization had already begun by 2 weeks (Fig. 2B & C & Fig. 3A), potentially stiffening those defects which were on a favorable progression toward regeneration.

An alternative but plausible interpretation of the data was that animals that were not recovering well from surgery were consequently not loading their operated hindlimb, and their poor prognosis for recovery was instigating lower strain amplitudes. However, this hypothesis was not supported, as initial gait analysis metrics representing operated hindlimb recruitment during walking revealed no relationships with bone volume or failure torque (Fig. S5D-I). Additionally, all animals recovered well from surgery with no visible signs of post-operative distress. These results suggest that early stage dynamic mechanical stimuli may play a persistent role in the healing progression and have the potential to influence tissue repair at later-stage time points.

### Strain magnitudes correlated with gait function and healing status

Another potential beneficial application for the sensor platform is to provide a non-invasive readout of the progression of healing. We hypothesized that the magnitude of fixation plate deformation would primarily be dictated by three factors: the force input on the operated femur during each step cycle, and the apparent stiffness of both the fixator and the adjacent bone defect, which share load as parallel deformable bodies. To obtain an indirect estimate of the hind limb force, we assessed the degree of operated hind limb utilization via weekly quantitative gait analysis immediately after treadmill rehabilitation periods. We observed significant deficiencies in mean paw print area and duty cycle of the operated hindlimb relative to the naïve contralateral initially after surgery, with no relative effects of fixator stiffness (Fig. 4A & B). These deficits were gradually restored to pre-operative levels within 6 weeks.

**Fig. 4:**
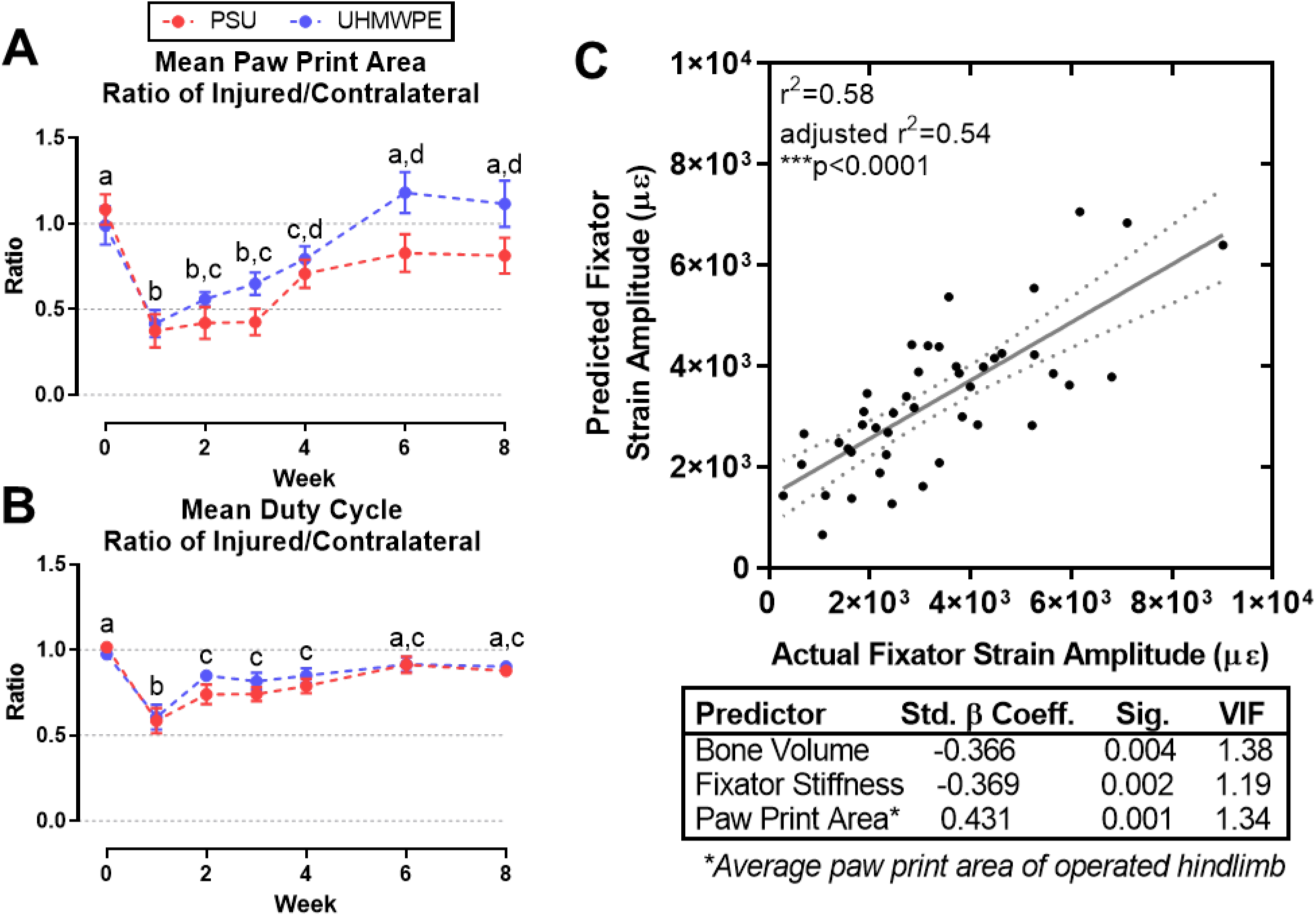
Strain magnitudes correlated with gait function and healing status. Longitudinal gait analysis quantifications for ratio of operated hindlimb over naïve contralateral for mean **(A)** paw print area **(B)** and duty cycle demonstrating substantial gait deficits are present 1 week after surgery and are progressively resolved by 6 weeks. n = 10-11. Differing letters denote significant differences p < 0.05 via Two-way ANOVA with Tukey’s test. **(C)** Regression model of pooled ambulatory strain measurements across all imaging time points demonstrate strain magnitude is predicted by a linear combination of bone volume, fixator stiffness, and mean paw print area. n = 44. Adjusted r^2^ = 0.54, ***p < 0.0001 via multiple regression.

To evaluate the accuracy of our simplified working model for fixator strain magnitude, we performed forward selection and backward elimination multivariate linear regression on a response variable data set consisting of all strain measurements acquired at time points with corresponding microCT scans (ranging from 2-8 weeks). The predictor variable pool consisted of the fixator stiffness of each device, bone volume, and gait analysis metrics. The optimal predictive model was identical for both directions and highly significant. Lending support to the working model, experimental inputs predictive of fixator strain magnitude consisted of negative correlations with bone volume and fixator stiffness, and a positive correlation with average paw print area of the operated hind limb (Fig. 4C). The three predictor variables exhibited low collinearity (VIF ≤ 1.38) and a maximal adjusted coefficient of determination overall, signifying parsimony. Together, these data help to describe the key biomechanical relationships within the long bone healing environment and demonstrate the potential for strain-based readouts to provide real-time assessment of mineralization progress during controlled functional activities without the use of X-rays.

### Defect revascularization at 3 weeks was elevated by load-shielding stiff fixation

Vascular perfusion of tissue is considered a critical precursor to osteogenesis. We investigated whether the mechanical environment conferred by UHMWPE also differentially affected vascularization. To this end, we performed a second in vivo study quantifying the size distribution of vascular structures using microCT angiography at 3 weeks, an intermediate time point that approximately coincides with bridging for the majority of healing defects (Fig. 5A-C). In vivo strain sensor measurements during gait replicated similar amplitudes and temporal trends as the preceding bone repair study (Fig. 5J and Fig. 2A), indicating the mechanical environment produced by rehabilitative walking was repeatable across studies. Regardless of fixator stiffness, we observed a significant increase in the amount of relatively small blood vessels (30-120 µm diameter) and the connectivity of the vascular network throughout the defect and surrounding tissue, demonstrating a potent angiogenic sprouting response to the injury and treatment with BMP-2 (Fig. 5D & H). Neovessel orientation was significantly more isotropic compared to naïve vasculature, which was primarily aligned along the limb axis (Fig. 5A-C, I).

**Fig. 5:**
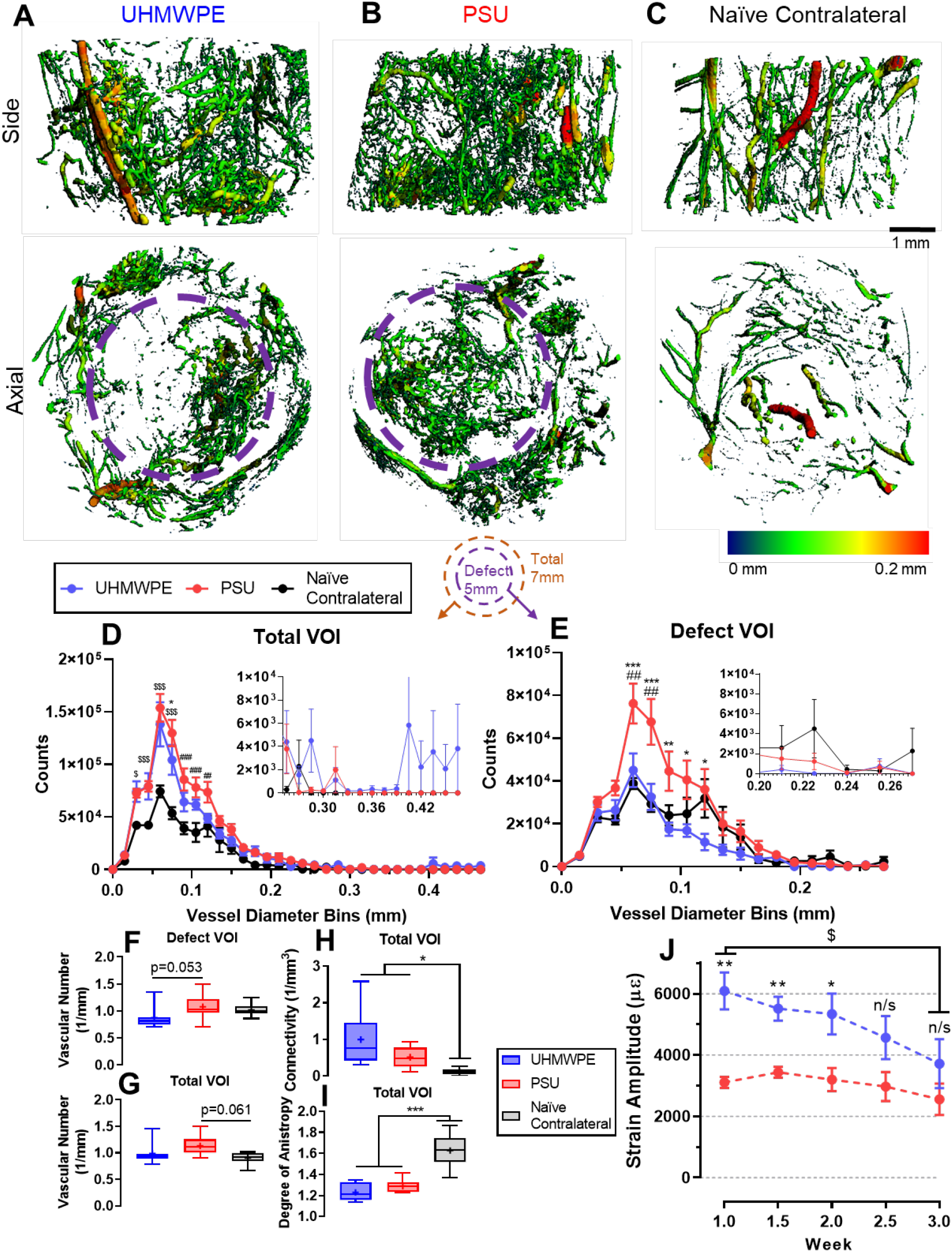
Defect revascularization at 3 weeks was elevated by load-shielding stiff fixation. Representative microCT reconstructions of blood vessels in the bone defect region at 3 weeks illustrate robust vascularization in operated hindlimbs stabilized by **(A)** UHMWPE and **(B)** PSU fixators relative to **(C)** naïve contralateral femora. Violet dashed circles delineate the location of the PCL tube, the interior of which is the Defect VOI. **(D)** Vascular thickness histograms demonstrate a significant increase in the presence of 45-90 µm thick blood vessels throughout the defect and surrounding tissue in operated femora, regardless of fixator stiffness, demonstrating pronounced angiogenesis in peripheral tissues in response to injury. **(E)** 60-120 µm thick blood vessels were significantly higher within defects stabilized by PSU fixators relative to the more compliant UHMWPE fixators. n=6-10. *p<0.05: PSU vs. UHMWPE, #p<0.05: PSU vs. naïve contralateral, $p<0.05: PSU & UHMWPE vs. naïve contralateral via Two-way ANOVA with Bonferroni pairwise comparisons. Vascular number in the **(F)** defect and **(G)** total VOIs, and **(H)** connectivity and **(I)** degree of anisotropy in the total VOI. n=6-10. *p<0.05 via Kruskal-Wallis with Dunn’s test. **(J)** Longitudinal in vivo strain amplitudes replicated the preceding 8 week study, with an initial two-fold increase in deformation on UHMWPE fixators which steadily declined, converging with PSU strain amplitudes. n=7-10. Two-way ANOVA *p<0.05 UHMWPE vs. PSU with Tukey’s test, $p<0.05 UHMWPE: week 1 vs 3 with Sidak’s test.

Interestingly, when we localized the analysis within the confines of the defect we observed a significant increase in the number of relatively small vessels (60-120 µm diameter) within defects stabilized by PSU fixators relative to more compliant UHMWPE fixators (Fig. 5E). The vessel size distribution of defects stabilized by UHMWPE more closely matched naïve vasculature. Furthermore, the number of distinct vascular structures within defects stabilized by PSU fixators was elevated (Fig. 5F & G). Together, these results suggest that angiogenic sprouting of new vessels within the defect is increased at 3 weeks due to low strain magnitudes under PSU fixation. More generally, the results demonstrate the utility of the sensor platform to quantify in vivo mechanical cues that exert potent but divergent magnitude-dependent effects on the formation of bone and vascular structures, respectively.

### Tissue-level compressive strains at 2 weeks were significantly elevated with compliant fixation

While the strain sensor provides a quantitative readout of axial strain on the fixator stabilizing the defect, the effects of mechanical stimulation are ultimately mediated by the tissue-level biophysical environment within the regenerative niche. To investigate this, we used the experimental data to inform image-based computational models of the early-stage bone defect mechanical environment. In this experimental system, the deformation measured on the fixation plate during gait provided an experimentally validated in vivo boundary condition for the fixator-femur system at 2 weeks (Fig. 6A-B). Using sample-specific boundary conditions and microCT geometry of the defect tissue, we observed a dramatic increase in the magnitude of the 3^rd^ principal strain, the maximum compressive local strain, within the defect soft tissue under compliant UHMWPE fixation (Fig. 6C-D). The defect tissue was largely load-shielded by PSU fixation, with 85% of the tissue undergoing less than 0.5% compressive strain (Fig. 6E). Conversely, load sharing permitted by UHMWPE fixation produced a much wider distribution of local tissue strain encompassing larger compressive magnitudes up to 6.7%. The average strain magnitude under UHMWPE was substantially elevated compared to PSU, and the interquartile ranges scarcely overlapped (Fig. 6F; UHMWPE: −0.3% to −3.0% vs. PSU: −0.1% to −0.4%). These data further demonstrate the utility of the implantable strain sensor platform to identify general tissue-level dynamic strain magnitudes that promote bone formation before appreciable mineralized bridging has occurred.

**Fig. 6:**
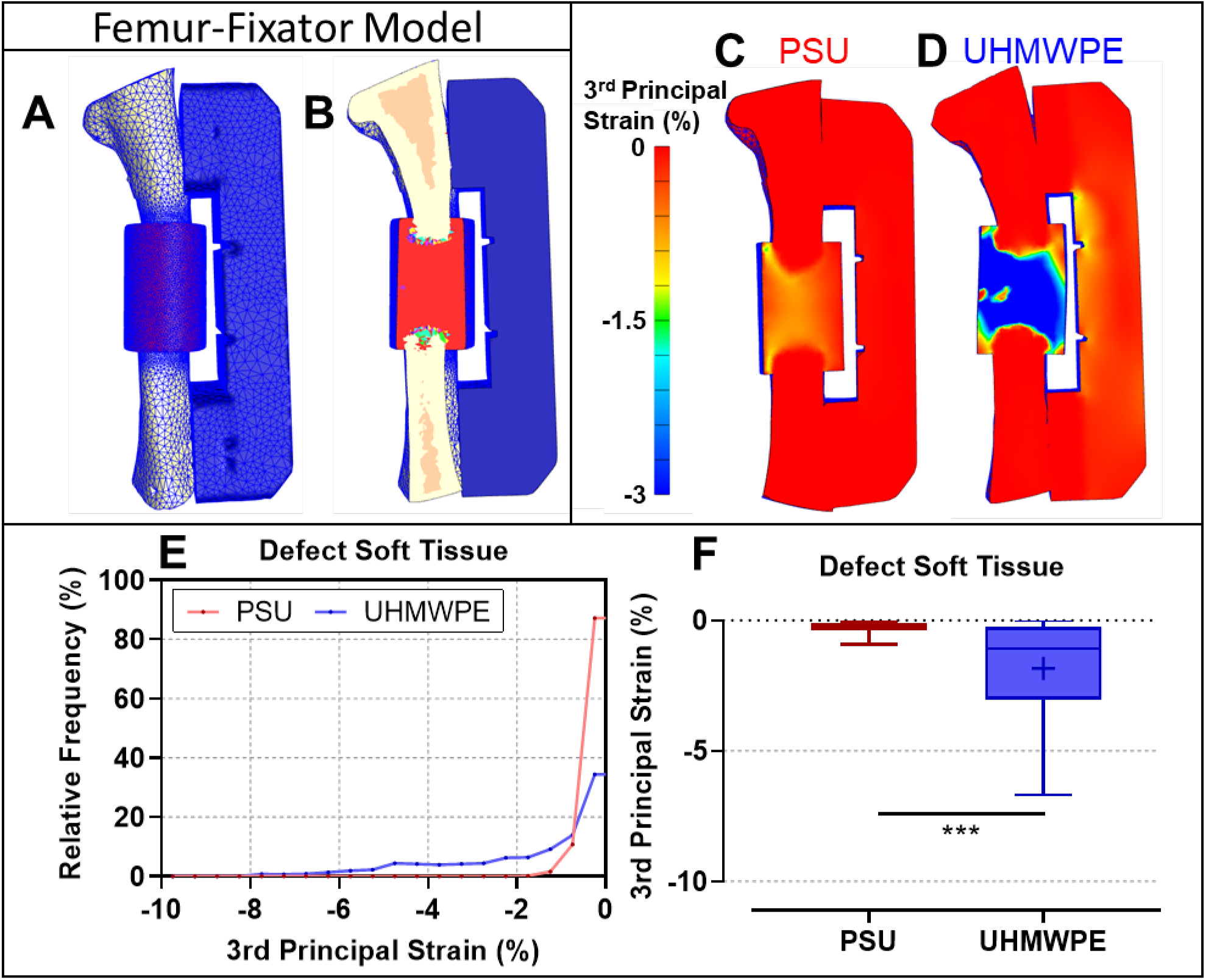
Tissue-level compressive strains at 2 weeks were significantly elevated with compliant fixation. **(A)** To assess the early-stage tissue-level mechanical environment produced by the differing fixators, microCT-based finite element models of representative samples stabilized by UHMWPE and PSU at 2 weeks were developed. Specimen-specific boundary conditions were validated using in vivo strain sensor measurements. **(B)** Cross-sectional view of finite element model, brightly colored elements in the defect zone represent immature woven bone. Local strain map of defects stabilized by **(C)** PSU **(D)** UHMWPE fixators reveal substantially elevated compressive strains throughout unmineralized tissue in the defect. **(E)** Histogram and **(F)** box plot further demonstrate significant increase in compressive local tissue strains permitted by compliant UHMWPE fixation. n > 16,000 elements with whiskers extending from the 2.5 to the 97.5 percentile values. ***p<0.001 via Mann-Whitney U test.

## Discussion

Physical forces critically regulate collective cell behavior during transient healing processes (*3*). However, dynamic forces are challenging to characterize in vivo, precluding a clear understanding of how external mechanical cues may be exploited therapeutically to enhance repair. Here, we sought to investigate the temporal progression of mechanical signals in a regenerating orthotopic site and to evaluate the biomechanical and therapeutic effects of a load sharing-inspired approach on bone regeneration. To this end, we integrated an implantable sensor platform into internal fixators of differing stiffness, creating an in vivo bone repair model in which we could perturb and remotely quantify mechanical boundary conditions in real-time during ambulation. We hypothesized that moderately compliant UHMWPE internal fixators would permit increased strain across the defect during gait relative to stiffer PSU and consequently enhance bone repair in a challenging critically-sized long bone defect small animal model. The results showed that load-sharing permitted by UHMWPE fixators initially delivered a two-fold increase in deformation magnitude, subsequently increased mineralized bridging by nearly three-fold, and increased bone formation by over 60% (Fig. 2A-C & Fig. 3A-B). Furthermore, by coupling in vivo imaging with strain sensor data as experimentally validated in vivo boundary conditions, we quantified differences in the early stage tissue-level mechanical environment mediated by reduced fixator stiffness (Fig. 6). Together, these results establish the utility of in vivo telemetric sensing approaches to aid in the measurement of tissue healing and outcomes after regenerative therapeutics. Additionally, the present study demonstrates that early stage physical stimuli exert potent effects on these healing outcomes and that the judicious use of rehabilitative loading activities informed by real-time biomechanical measurements may have the potential to resolve skeletal injuries more effectively.

At the tissue level, the benefits of mechanical stimulation to skeletal repair have been largely established using percutaneous and bulky external fixator frames; with these systems loads are applied via manual adjustment, actuators, or spring systems, but the invasiveness of such approaches substantially limits their utilization (*8*, *27*). At the cellular level, mechanical stimulation is known to differentially regulate a wide array of mechanisms critical to tissue regeneration including mesenchymal stromal cell differentiation and migration (*2*, *28*) and vascular formation and remodeling (*17*, *29*). Together, data from large and small animal models implicate certain mechanical cues may enhance bone repair in a strain magnitude and healing stage dependent manner (*6*). Invasive external loading systems applying non-physiological loading patterns have indicated that relatively small early compressive strains (3-7%) support early callus precursor formation and woven bone formation (*8*, *30*). Early-stage deformation exceeding 10-15% has been implicated to inhibit vascularization and cause fibrotic non-union or fracture the newly formed bone bridge, though in vivo strains were not actually measured (*17*, *31*). The defect stiffens by several orders of magnitude as mineralization and bridging occurs allowing larger magnitude loads to be exerted while resultant tissue strains decrease; osteogenic strain magnitudes on newly formed woven are well established to be approximately 0.1-0.4% and can accelerate mineralization and vascular maturation after the interfragmentary gap has already bridged with calcified tissue (*7*, *17*, *25*, *32*, *33*). Taken together, the mechanobiological regulation of bone (re)modeling after bridging is well established. However, the magnitude of and functional effects of physiological mechanical cues imparted by rehabilitative activity like walking during early phases, prior to the establishment of bridging or non-union, remains poorly understood. Clinically, it would be beneficial to leverage the potential of local mechanical therapies via early loading, since later stage interventions offer limited benefit to vulnerable cases at risk of non-union. Due to the lack of understanding of early loading and technologies to quantify it in vivo, current rehabilitation protocols for fractures treated with open reduction and internal fixation are typically non-weight bearing for 6-12 weeks (*34*).

An alternative mechanobiological strategy is to introduce mechanical cues into the regenerative niche non-invasively via ambulatory activities. In addition to direct biophysical stimulation of the skeletal defect, rehabilitative exercise may exert second-order benefits to peripheral tissues injured during trauma or surgery including surrounding muscles and nerves (*15*, *16*). Randomized clinical trial evidence has shown that supervised progressive resistance exercise after proximal hip fracture enhances physical function and quality of life compared to low-intensity exercise targeting flexibility for elderly patients (*14*). However, the actual mechanical environment imparted by rehabilitative activity, such as resistance training, within a bone defect has not been quantified longitudinally, impeding a generalized understanding of tissue-level mechanical stimuli with the potential to augment regeneration. The in vivo experimental system that we developed and used in the studies herein allowed us to longitudinally measure dynamic axial strain during gait. Concurrently, we observed a substantial enhancement in bone repair via increased load sharing. These data implicate a critical role for early mechanical cues on the long term healing response as strain cycle magnitude at 1 week (before appreciable healing occurred) had a significant positive correlation with the long-term bone regeneration outcomes (Fig. 5A-C).

While fixator strain measurements were useful to assess differences between groups and temporal trends reflective of healing, mechanical cues transmitted to cells within the healing tissue ultimately regulate mechanobiological responses. Given the potent longer-term osteogenic effects of early-stage loading under compliant UHMWPE fixation, we reasoned that quantifying the tissue-level deformation provided a more generalizable understanding of dynamic mechanical cues that accelerate early-stage bone repair, irrespective of the fixator. Subject-specific, image-based finite element analysis revealed that tissue-level compressive strain within the bone defect at 2 weeks ranged from 0.1-6.7% under UHMWPE fixation, whereas stiffer PSU fixation shielded the entire defect, permitting only 0.1-0.9% strain. These data are consistent with the results of Miller et al., who reported that local maximum compressive strains of about 3% resulted in peak probability of subsequent mineralization between days 7 and 14 using an invasive percutaneous loading system in a rat osteotomy model (*30*). Miller and colleagues further showed that the probability of fibrous tissue formation surpasses mineralization when compressive strains exceed 8%. The results from this study corroborate the osteogenic nature of early-stage tissue strains below 7% using a completely implantable telemetric system and leveraging ambulatory activity to non-invasively deliver mechanical cues. Together, the data reported here support the hypothesis that mechanobiological responses to ambulatory loading can be therapeutically exploited to accelerate bone repair, and further research is warranted to investigate how controlled rehabilitation regimens may be safely enacted relatively early after surgical intervention.

Angiography measurements revealed that the beneficial effects of load sharing on bone formation differentially regulated defect revascularization at 3 weeks post-surgery. Our previous work in this model used a compliant fixator possessing a stiffness about 50% lower than the UHMWPE fixators; the former fixators with lower stiffness dramatically impaired both vascular ingrowth and bone regeneration after injury (*17*). However, in the current study we observed a potent osteogenic effect from the deformation permitted by UHMWPE fixation possessing a more moderate compliance. Interestingly, load shielding under PSU fixation increased the number of relatively small vessels in the defect at 3 weeks, whereas UHMWPE fixation reached similar vessel number and size distribution as the naïve contralateral femur. Together with our previous work, these data suggest a potential minimum threshold of vascularization is necessary to support osteogenesis, above which there may not be further benefit to the regenerative capacity of the tissue. Overall, there appear to be distinct magnitude-dependent mechanobiological thresholds that differentially impair either bone or neovascular growth when exceeded. As a result, increased strain magnitude can slightly modulate the progression of angiogenesis but still support sufficient tissue revascularization to significantly enhance bone repair. These results contrast the traditional paradigm in which increased angiogenesis correlates with enhanced osteogenesis; and instead, implicate a minimum threshold of angiogenesis after which point there is not necessarily an additive therapeutic benefit. Further, these data show that this threshold is highly mechanosensitive and likely varies temporally during different stages of regeneration.

In addition to the mechanobiological findings facilitated by the strain sensor platform, the data support that future iterations of strain sensing approaches may have promising applications in clinically relevant contexts such as diagnostics and rehabilitative monitoring. While not the primary impetus for the sensor platform developed in this model, the results provide proof of principle as strain amplitudes during gait gradually declined over time in healing defects. The magnitude of a given strain measurement was a function of the volume of mineralized tissue in the defect, the degree of hindlimb usage, and the fixator stiffness (Fig. 4C). Therefore, this study provides pre-clinical in vivo evidence that it may be feasible to infer the status of bone healing via measurements acquired under a repetitive controlled task without the need for X-rays. Such an approach is pertinent to pediatric patients after procedures that would typically entail CT imaging, as radiation exposure should be curtailed to minimize risk of radiation-induced cancers (*35*). This idea has been explored of late in percutaneous devices to measure strain and tissue impedance (*36*, *37*), but emerging developments in passive and active devices could offer fully implantable, wireless embodiments (*38*). In addition, we obtained measurements in real-time while animals walked on a treadmill. Such an approach could provide real-time feedback to guide rehabilitative specialists while patients perform functional activities, enabling patients to maximize recovery of function after surgery while ensuring hardware safety thresholds are not exceeded.

The sensor platform does have limitations that warrant further research. Though sufficient for implantation in a relatively small rat model, the size of the fully-packaged transceiver (38 mm × 23 × 12 mm) would benefit from miniaturization. The primary contributor to the size was the coin-cell battery, which was selected for its relatively high power density and sufficient theoretical power budget for an 8 week study. Nonetheless, transceiver attrition was unexpectedly high in the initial bone repair study, reaching 33% and 83% by 3 and 8 weeks, respectively. Explant testing indicated this was due to variability in voltage regulator performance. Improvements to the transceiver voltage regulator in the follow-up angiography study substantially increased device reliability. Attrition was reduced to a single failed device (6% failure rate) at 3 weeks, where the root cause was due to fixator loosening and not electrical malfunction. Additional work toward circuit board layout and power consumption optimization would substantially reduce device size and improve power budget fidelity. Passive approaches for power input and data transfer are a promising avenue for sensor miniaturization in certain contexts, but require externally mounted inductive coil elements, have reduced transmission range, and are sensitive to variation in coil-pair orientations. Together, these are significant limitations to monitoring during dynamic movements. For these reasons, we employed an active BLE telemetry approach that permitted straightforward sleep-wake control and parallel data acquisition from multiple animals walking at once. Despite the size of the transceiver, it is worth noting that the current footprint is than sufficient for large animal and human scale hardware research.

The specific objective of this study was to assess the role of the tissue-level mechanical environment on regeneration. Treadmill walking at a constant speed was employed as a relatively controlled ambulatory activity with which to impart mechanical stimulation for all groups. Therefore, the net contribution to tissue repair of periodic walking relative to cage restriction was not targeted, but warrants further investigation. Forced walking may cause stress and confounding systemic effects in small animals (*16*). The findings of this study motivate the development of experimental rehabilitative protocols that better mimic clinical regimens in both small and large animal models to improve translation of insights obtained by in vivo strain sensing platforms to humans, similar to the methods of Dalise and colleagues studying aerobic exercise in rats (*39*).

Here, we implemented an implantable sensor platform to monitor how mechanical stimulation of the regenerative niche permitted by reduced stiffness fixation promotes bone repair. In this study, in vivo strain monitoring enabled observations that: 1) early strain amplitudes correlated with healing outcomes before radiographic indications of healing were apparent, and that

2) local strain magnitudes within the regenerative nice between 1-7% significantly enhanced bone regeneration. In contrast to the generally accepted paradigm directly linking the enhancement of angiogenesis with improved osteogenesis, this study indicated that mechanical stimulation can slightly reduce early vascular ingrowth but still support adequate revascularization to significantly augment bone regeneration. These data motivate further inquiry into mechanosensitive thresholds which are critical regulators of both angiogenesis and osteogenesis and have broad implications for regenerative medicine. By facilitating real-time longitudinal characterization of in vivo mechanical cues, the data represent a notable advancement in regenerative mechanobiology. The capacity of integrated biomedical sensors to remotely quantify dynamic mechanical signals throughout musculoskeletal regeneration offer new opportunities to investigate mechanisms by which biophysical cues augment tissue repair. With continued research, similar sensor approaches have the potential to provide instantaneous clinical feedback for physicians and physical therapists to aid implementation of safe and effective rehabilitation regimens optimized for restoration of tissue structure and function.

## Methods

### Device fabrication

A new digital transceiver unit was developed to provide a number of key functional improvements over the unit reportedly previously (*22*), mainly: (i) increased power budget and (ii) sampling frequency, (iii) improved connectivity via Bluetooth Low-Energy (BLE) wireless network, and (iv) remote-controlled circuit calibration. The unit used a BLE microcontroller (MCU) (Silicon Labs BLE113) to receive commands and transmit data at an increased frequency of 30 Hz through a PC-mounted USB receiver. A 620 mAh battery (Panasonic CR2450) was used to provide an 8 week power budget. Furthermore, a low-power receiving “sleep” mode was implemented, allowing the PC user to remotely activate to the unit to acquire and transmit measurements as needed, permitting flexible measurement time points and durations. A wirelessly-controlled digital rheostat was also implemented to allow remote calibration of the baseline voltage signal while implanted. The transceiver was encapsulated in a custom 3-d printed housing (Form 2, Formlabs) fully encapsulated in an ISO 10993 compliant UV-curing urethane (Dymax).

As previously described (*22*), the transceiver was connected via two 36 AWG braided stainless steel wires encapsulated in biocompatible silicon tubing (AM systems) to a single-element 350 Ω strain sensor (PGEA-06-125BZ-350/E, Vishay) integrated into a custom internal fixator used to stabilize the femoral defect. To modulate fixator stiffness and load sharing across the defect, the radiolucent, polymeric bridging element was either polysulfone (PSU, McMaster-Carr) or ultra-high molecular weight polyethylene (UHMWPE, Quadrant), creating “stiff” and “compliant” fixator groups, respectively. Prior to implantation, each device was calibrated in three-point bending (TA Electroforce 3220) to strains encompassing physiological magnitudes.

### Surgical procedure

Identical procedures were used in both bone repair and angiography studies. As previously described (*22*), a unilateral critically-sized 6 mm segmental defect was created in the left femur of 15-wk-old female CD (Sprague-Dawley) rats (Charles River Labs). Femurs were stabilized by either stiff (PSU) or compliant (UHMWPE) fixators with integrated strains sensors. Transceiver packs were mounted in the abdominal cavity and connected via a tunneled lead passing through a keyhole incision in the abdominal wall superior to the left inguinal ligament into the hindlimb compartment. Defects were either treated with 2 µg recombinant human bone morphogenetic protein 2 (BMP-2, Pfizer), delivered via a hybrid biomaterial scaffold described below consisting of 120 µL BMP-2-laden alginate hydrogel injected inside an electrospun polycaprolactone (PCL) tube, or left empty. Empty defects form negligible mineralized or soft tissue and served as non-healing negative controls. Animals were randomly allocated to experimental groups. Animals were anesthetized and then euthanized by CO_2_ asphyxiation at either 3 or 8 weeks. All procedures were approved by the Georgia Institute of Technology IACUC (Protocol A17034).

### Hybrid RGD-alginate/PCL scaffold production

Detailed scaffold production methods are described elsewhere (*40*, *41*). Briefly, RGD-functionalized alginate (FMC BioPolymer) was reconstituted in α-MEM (Thermo Fisher Scientific) at 2% w/v. Recombinant human BMP-2 (Pfizer) was reconstituted in a 0.1% solution of rat serum albumin (Sigma-Aldrich) and 4 mM HCl and mixed with alginate, yielding 2 µg BMP-2 per 120 µL. The solution was ionically cross-linked with a 0.21 w/v CaSO_4_ slurry mixed at a 1:25 volume ratio. Sheets of PCL nanofiber mesh were generated by electrospinning a 12% w/v solution of PCL in 90:10 (v/v) of hexafluoro-2-propanol:dimethylformamide from a syringe at a flow rate of 0.75 mL/h and a 15-20 kV potential onto a grounded collector plate 20-23 cm away. PCL sheets were laser cut into rectangles with a chessboard grid of 23 x 1 mm circular perforations, rolled and glued into 5 mm cylinders using ISO 10993 UV-curing adhesive (Dymax).

### Gait analysis

Gait capture was performed longitudinally to assess hindlimb utilization for each rat using a Catwalk 7.1 system (Noldus). Rats were placed on an illuminated runway and allowed to walk freely between either ends. Illuminated paw prints were recorded by a digital camera and runs where the rat traversed the entire length of the runway were analyzed (2-3 runs per animal, 10 second max) before surgery for baseline, and 1, 2, 3, 4, 6, and 8 weeks after surgery. Automated footprint classifications were verified and corrected manually for each run. Paw print area and duty cycle (ratio of stance duration to the sum of stance and swing duration) were computed.

### Treadmill walking

Before surgery, rats were acclimated to walk consistently on a treadmill (NordicTrack) at speeds ranging from 5-9 m/min over a period of 10-15 min. Beginning 1 week after surgery, and twice weekly thereafter, each animal was walked for 10 min at a consistent speed of 6.5 m/min, creating a gait cycle of approximately 1-2 Hz. The total distance travelled loosely approximated the distance traversed during one day of in-cage activity (*42*).

### Strain measurement and analysis

During treadmill walking sessions, strain measurements were transmitted in real-time via Bluetooth to a nearby laptop and plotted on a custom Visual Studio C# (Microsoft) graphical user interface (GUI). Data was collected for 3 minutes of the treadmill session. After 4 weeks, strain was measured during the first treadmill period of the week. A custom MATLAB (Mathworks) script was developed to identify local maxima and minima pairs in the signal corresponding to individual step cycles, and strain amplitudes were computed.

### X-ray video collection

One month after surgery, high-frame rate images of an empty defect control animal’s skeleton were obtained to demonstrate sensor functionality during treadmill data collection. A custom radiographic imaging system comprised of bi-plane X-ray generators, image intensifiers (Imaging Systems & Service, Inc.), and high-speed digital video cameras (Xcitex XC-2M, Woburn, MA) was used to collect 6 second videos (100 frames/s; 43 kV, 100 mA, 5 ms exposure).

### In vivo radiographs and microCT

In the bone repair study, in vivo radiographs and microCT scans were acquired at 2, 4, and 8 weeks to assess bone formation and bridging across the defect. Digital radiographs were acquired at 25 kV with a 15 sec exposure (Faxitron MX-20). MicroCT scans (VivaCT 40, Scanco Medical) were performed using 26.3 µm voxels, 55 kVp, 145 µA, and 300 ms integration time. Scans were manually aligned along the femoral axis prior to analysis. Mineral formation was evaluated inside a cylindrical volume of interest (VOI) of 5 mm in diameter and 4 mm long centered between the intact bone ends and encompassing the defect and PCL electrospun tube. Polar moment of inertia was assessed with a VOI encompassing all mineralized tissue throughout the entire defect and 1 mm of each bone end to include periosteal mineralization at the defect boundaries. Bone was segmented by applying a Gaussian filter (sigma=1.2, support=1) and a global threshold of 388 mg hydroxyapatite/cm^3^, corresponding to half the density of intact cortical bone.

### MicroCT angiography

Vascular perfusions were performed after 3 weeks in the angiography study (*43*). Animal vasculature were sequentially perfused through the ascending aorta with 0.9% saline, 0.4% papaverine hydrochloride vasodilator, 0.9% saline, 10% neutral buffered formalin (NBF), 0.9% saline, and radiopaque lead chromate contrast agent (2:1; Microfil MV-22, FlowTech Inc.). Samples were stored overnight at 4° C to ensure polymerization, and both operated and naïve femora were dissected with surrounding musculature left intact. Samples were submerged in a formic acid/citrate decalcifying solution (Newcomers Supply) for 10 days on a rocker plate with daily solution changes.

MicroCT scans were performed using 15 µm voxels, 55 kVp, 145 µA, and 300 ms integration time. Vascular formation and morphology was assessed inside two different 4.14 mm long cylindrical VOI: a 5 mm diameter “Defect VOI” encompassing the interior of the PCL tube in operated femora or the approximate central axis of the femur in naïve contralateral samples, and a 7 mm diameter “Total VOI” encompassing the bone defect and immediate surrounding tissue. Vasculature was segmented using a Gaussian low-pass filter and a global threshold.

### Biomechanical testing

After eight weeks, animals were euthanized by CO_2_ asphyxiation, hindlimbs were cleaned of soft tissue, fixators were carefully removed, and femora were wrapped in PBS-soaked gauze and stored at −20° C until testing was performed 3 days later. Each femur was thawed before potting the ends in Wood’s metal (Alfa Aesar). Specimens were tested to failure in torsion at 3°/sec using a load frame (TA Electroforce 3220). Failure strength was defined as the peak torque over the first 40° of rotation, and stiffness was assessed as the slope of the linear region of the torque-rotation curve.

### Histology

One representative femur from both UHMWPE and PSU groups was selected for histology based on 8 week MicroCT bone volume. Samples were fixed in 10% NBF for 48 h at 4° C, then switched to PBS and shipped to HistoTox Labs for decalcification, paraffin processing, and staining. Midsagittal 5 µm thick sections were stained with H&E, Safranin-O/Fast Green, or Picro Sirius Red.

### Finite element analysis

Finite element simulations of the femur-fixator system at 2 weeks were generated for representative UHMWPE and PSU samples. Models were first created in MIMICS (Materialise) using animal- and time-specific in vivo microCT images. Image noise was suppressed using a Gaussian filter (sigma=1.2, support=1). The calcified tissue mask was cropped to only include the defect region and a sequence of erosion (radius =2) and dilation (radius=1) steps were applied to remove small volumes. Generalized microCT-based geometries of the proximal and distal femur segments were manually aligned to the animal-specific bone segments visible in the microCT images. Unmineralized portions of the defect region were assumed to be a homogeneous cylinder of soft tissue (consisting of soft callus tissue and alginate) with a diameter of 5 mm (the diameter of the PCL tube), span the length of the defect, and centered about intact distal and proximal femoral segments. Mineralized bone was subtracted from the soft tissue mask to ensure continuity between mineralized bone and soft tissue. The PCL tube mask was aligned with and enveloped the soft tissue mask. Fixator and steel riser plate models were manually aligned with the fixator plate visible in microCT images. All masks were defined separately for each material (soft tissue callus, mineralized tissue, proximal and distal bone segments, fixator risers and plate, PCL mesh) and were combined to establish one unified surface model. The unified surface mesh was then cleaned in 3Matic (Materialise) and converted to a volumetric mesh of quadratic 10-node tetrahedral elements. To determine mesh resolution, a convergence analysis was conducted using the 3^rd^ principal strain within the defect soft tissue as the convergence criterion (Fig. S6A).

A summary of material properties applied to the femur-fixator model is listed in Table S3. When available, material properties were assigned based on known properties (e.g., steel, cortical bone). The elastic moduli for the UHMWPE and PSU materials in the fixator were iteratively determined by matching model calculated reaction forces to those calculated during a three-point bending experiment (Fig. S6B-C). Similarly, elastic moduli for soft tissue callus (Fig. S6D) and mineralized tissue (Fig. S6E) were also matched to experimentally determined displacements. Briefly, femora with an unbridged defect (Fig. S6D) or a bridged defect (Fig. S6E) were excised, potted in Wood’s metal and tested in uniaxial compression with a ramp to 0.2 mm and 5.3 N, respectively. Soft callus tissue was assumed to be homogenous with a single elastic modulus, while elastic modulus of the mineralized tissue within the defect was assumed to be related to the density of independent voxels to a power of 1.49. A scalar factor was then multiplied to this relationship and adjusted to match the model-derived with the experiment-derived displacements. All materials were modeled as compressible neo-hookean with poisson’s ratios from the literature (*44*, *45*).

The boundary conditions assigned to the femur-fixator model were implemented to best reflect in vivo loading as determined by Wehner and colleagues (*46*). The distal femoral surface was fixed in all directions. Compressive and bending pressures, below, were applied to the cortical bone of the proximal surface (assumed to be 0.45 mm thick). Elemental pressures were formulated as a function of the medial-lateral and anterior-posterior position of each proximal surface element and a single scalar factor.

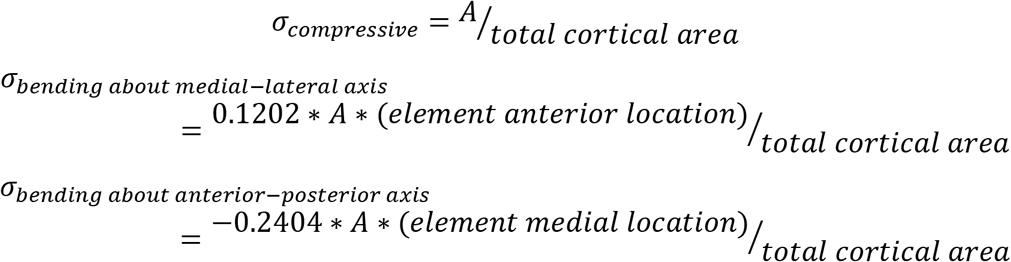

The scalar factor *A* was then iteratively modified until the average axial Lagrangian strain in the sensor region of the fixator matched the in vivo strain amplitude for the respective animal at 2 weeks. All finite element analyses were completed using FEBio (version 2.8.5) and analyzed in PostView (*47*).

### Statistical analysis

Sample sizes for treated groups were chosen based on a priori power analysis of prior mechanical loading model experiments. All statistical analyses were performed using Prism 7 (GraphPad), except multivariate linear regression, which was performed using SPSS 24 (IBM). Normality of data was tested using Shapiro-Wilk test. Fixation plate stiffness and sensor sensitivity were assessed by Student’s t-test and Pearson’s linear regression, respectively. Two-way ANOVA with Tukey’s or Sidak’s pairwise comparisons were used to compare groups for longitudinal strain, microCT, and gait analyses. Bonferroni comparisons were used to assess pairwise differences in vascular thickness distribution. Proportions of bridged defects were evaluated by chi-square test for trend, with pairwise comparisons assessed by individual chi-square tests. Non-parametric Mann-Whitney U or Kruskal-Wallis tests were used to evaluate non-normally distributed data torsion and vascular morphometric data. Spearman’s rank-order correlations were used to assess relationships between early strains and longer-term bone volume and strength. Data are displayed as mean ± s.e.m. or as box plots showing 25th and 75th percentiles, with whiskers extending to minimum and maximum values, unless otherwise noted in the figure heading.

## Supporting information

High-speed X-ray video of real-time strain acquisition during gait 1 month after surgery.

## General

We thank Dr. Albert Cheng, Dr. Ramesh Subbiah, Casey Vantucci, Ryan Akman, Lina Mancipe-Castro, Gilad Doron, Shannon Anderson, Joy Dutta, Fabrice Bernard, and Jay McKinney for assistance with animal surgeries. We acknowledge the Georgia Tech Physiological Research Laboratory and Petit Institute Core Facilities for their services and shared resources used in this work. We also thank Dr. Johnna Temenoff for rodent treadmill access, Dr. Young-Hui Chang and Nathan Kirkpatrick for access and assistance with high-speed x-ray video collection.

## Author contributions

B.S.K., N.J.W., and R.E.G. designed the research and performed surgeries; B.S.K. built devices and performed in vivo analyses; B.D.N, S.S.K., and K.G.O. designed and built transceivers; B.S.K., J.A.W, J.K., and S.J.H. performed finite element analyses; M.A.R. performed vascular perfusions; B.S.K and J.K. analyzed the data and wrote the paper; All authors edited and approved the final paper.

## Data and materials availability

All data needed to evaluate the conclusions in the paper are present in the paper and/or the Supplementary Materials. Additional data related to this paper may be requested from the authors

## Supplementary Materials

**Fig. S1:**
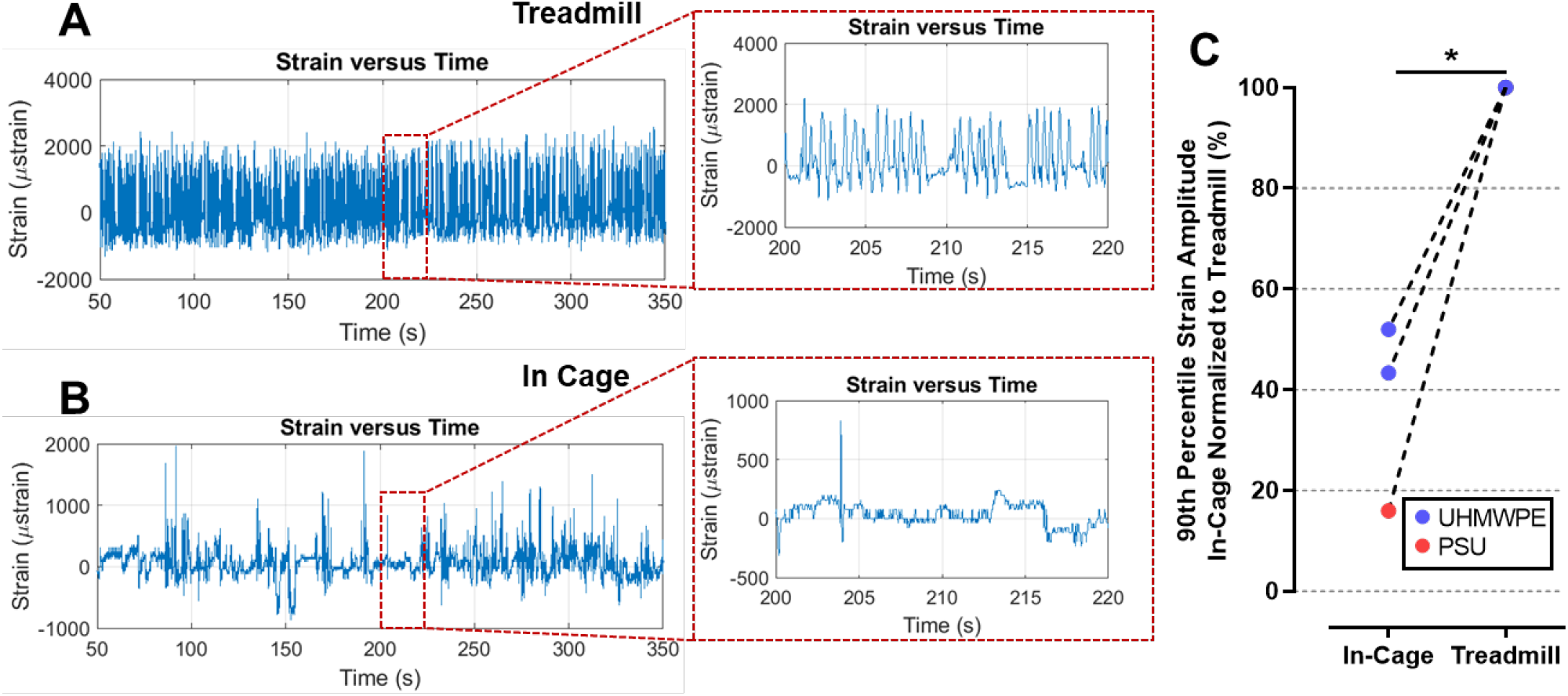
Treadmill walking is greater mechanical stimulus than in-cage activity in terms of both magnitude and frequency. Representative experimental strain measurements from the same animal acquired during **(A)** treadmill walking **(B)** and ad libitum nocturnal in-cage activity the same night. **(C)** 90^th^ percentile strain magnitudes during treadmill activities were 60% higher than corresponding nocturnal in-cage activities. n = 3. *p < 0.05 via paired t-test.

**Fig. S2:**
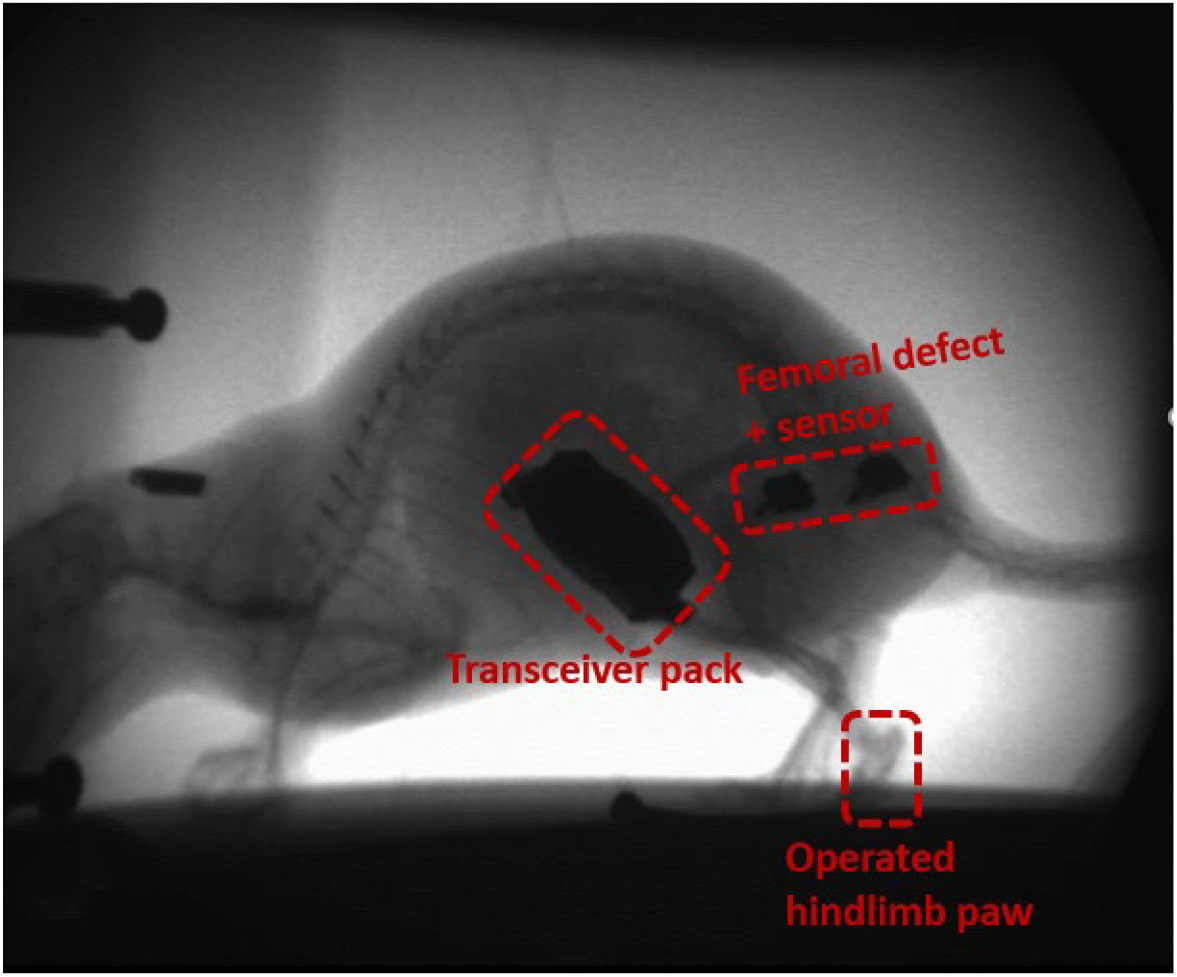
Annotated still image of high-speed X-ray video delineating femoral defect, corresponding hindlimb paw, and transceiver pack.

**Fig. S3:**
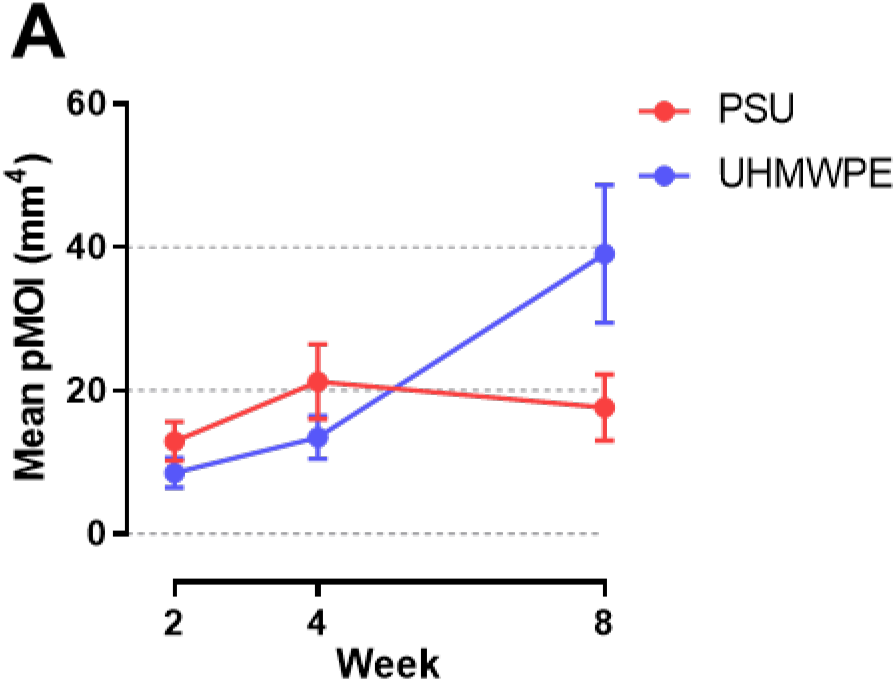
Longitudinal microCT quantification of additional morphometric parameters. **(A)** Mean polar moment of inertia was not significantly affected by fixator stiffness. n = 10-11. Vertically oriented bar **p < 0.01 overall main effect UHMWPE vs. PSU via Two-way ANOVA.

**Fig. S4:**
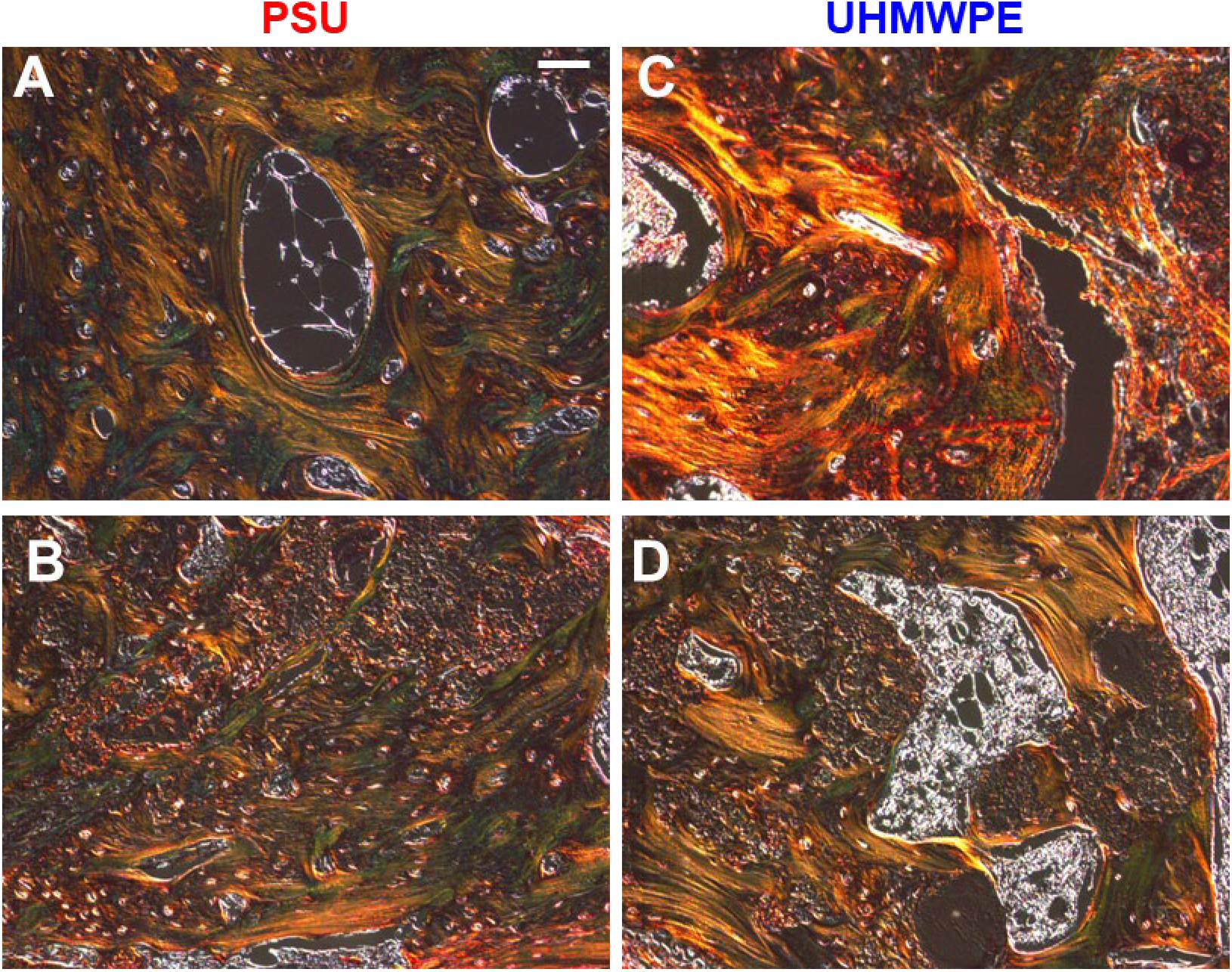
Extracellular matrix organization. Histological images of Picro Sirius Red stained sections at 8 weeks viewed under polarized light. **(A-B)** Defects stabilized by stiff PSU fixators exhibited primarily lamellar ECM possessing a mix of green and yellow/orange collagen fibers, indicative of a mixture of both smaller and larger fibers, respectively. **(C-D)** Defects stabilized by compliant UHMWPE fixators possessed a range of ECM organizations. **(C)** Large and intense yellow/orange collagen fibers were visible, indicative of newly formed woven bone and pronounced matrix remodeling. **(D)** In addition, regions with more organized green and yellow/orange fibers analogous to PSU samples were also apparent. Scale bar, 50 µm.

**Fig. S5:**
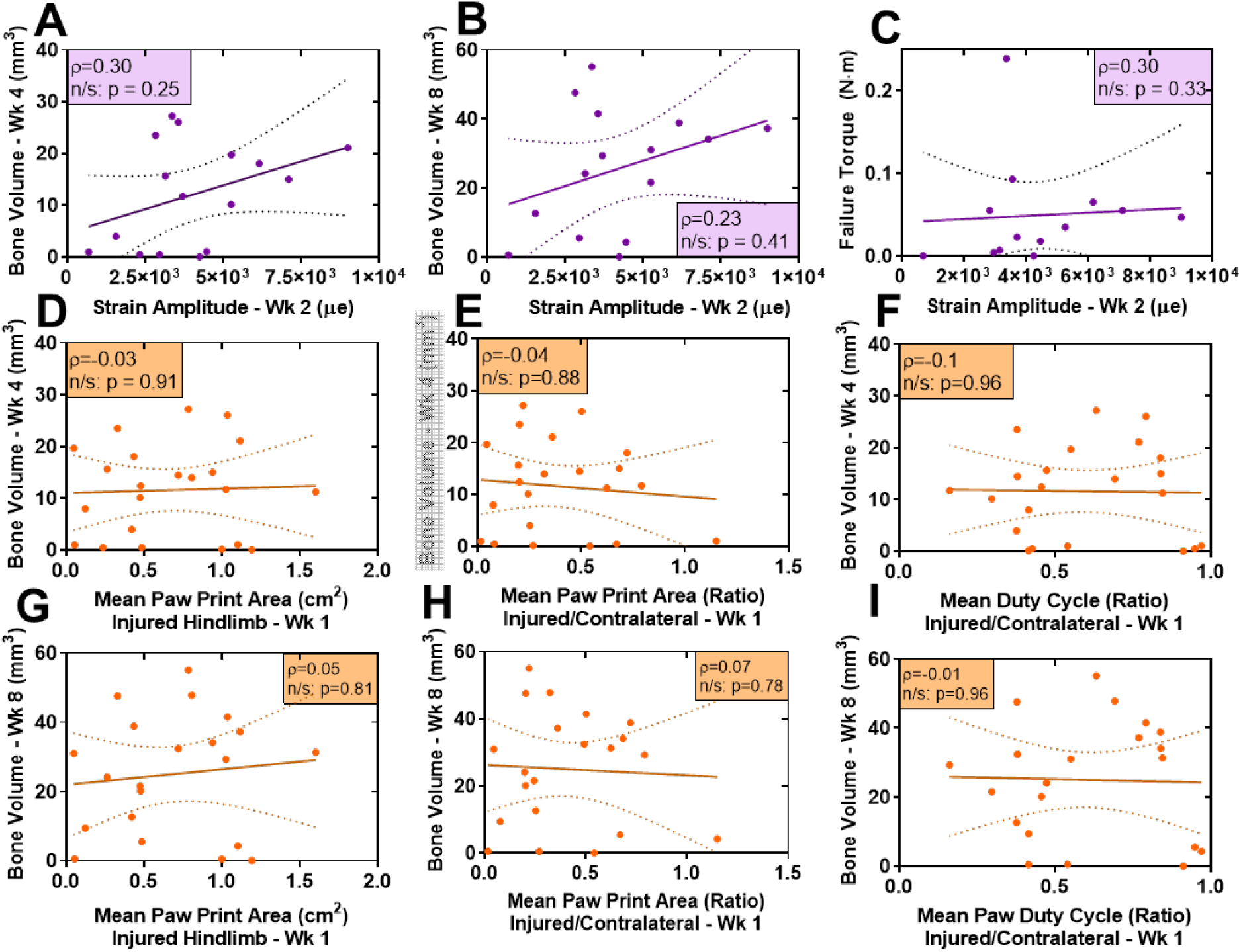
Later time point strain amplitudes and initial gait analysis metrics do not correlate with healing outcomes. Strain amplitudes acquired at 2 weeks no longer correlated significantly with **(A)** 4 week bone volume, **(B)** 8 week bone volume **(C)** and 8 week failure torque, as mineralization had initiated thereby stiffening defects with a favorable prognosis. n = 16. n/s via rank-order correlation. Gait analysis metrics at 1 week including mean paw print area of injured hindlimb and injured to contralateral ratios of mean paw print area and duty cycle also did not correlate with 4 week **(D-F)** or 8 week **(G-I)** bone volume, respectively, indicating that animals with increased gait deficits were not predisposed toward poor healing outcomes. n = 21-22. n/s via rank-order correlation.

**Fig. S6:**
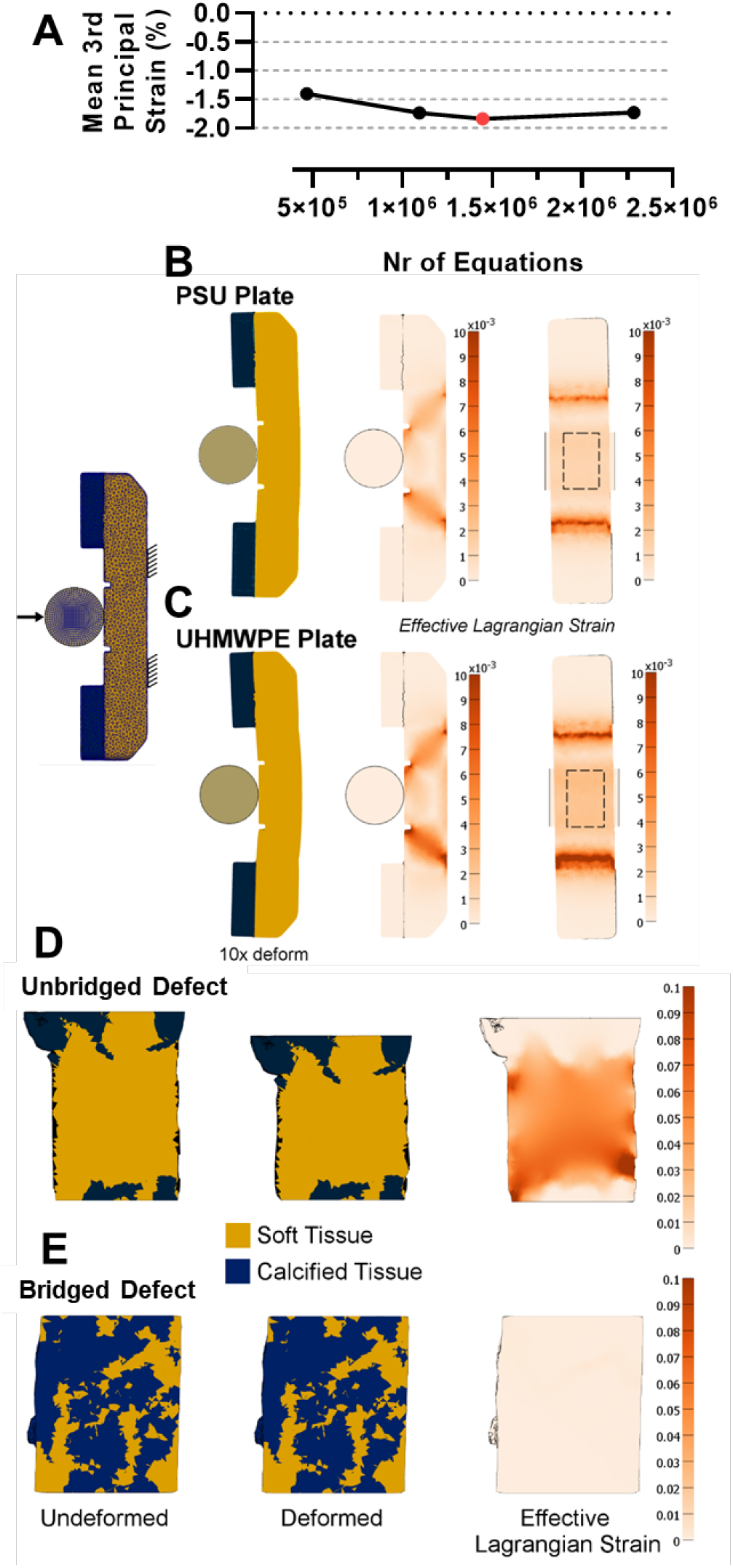
Finite element simulation development. **(A)** Femur-fixator finite element models were meshed using quadratic tetrahedral elements at increasingly refined resolutions until soft tissue 3rd principal strain converged. The red data point denotes the mesh resolution used here, where meshes consisted of approimately1.5M equations. Fixator mechanical properties were validated to match experimental three-point bending tests for both **(B)** PSU and **(C)** UHWMPE. Similarly, defect soft tissue and mineralized tissue mechanical properties were validated to match ex vivo experimental compression testing of **(D)** unbridged and **(E)** bridged defects.

**Table S1:**
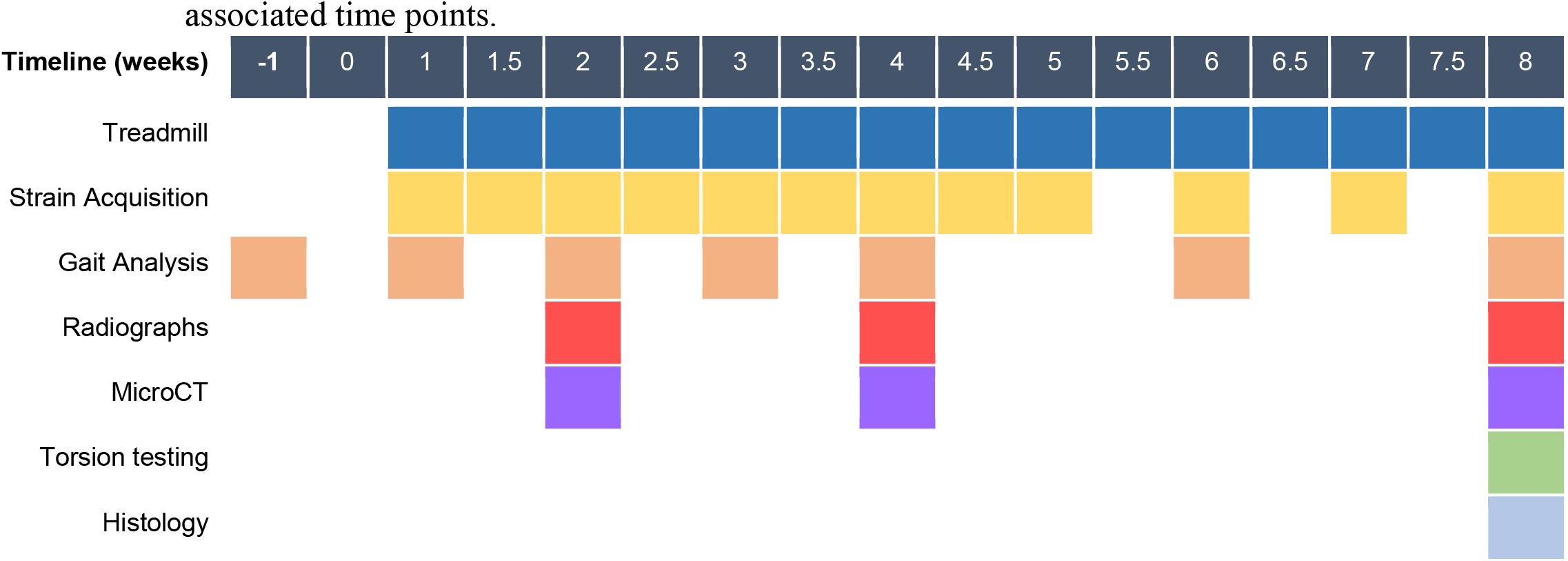
Bone repair in vivo study timeline. Experimental measurements and associated time points.

**Table S2:**
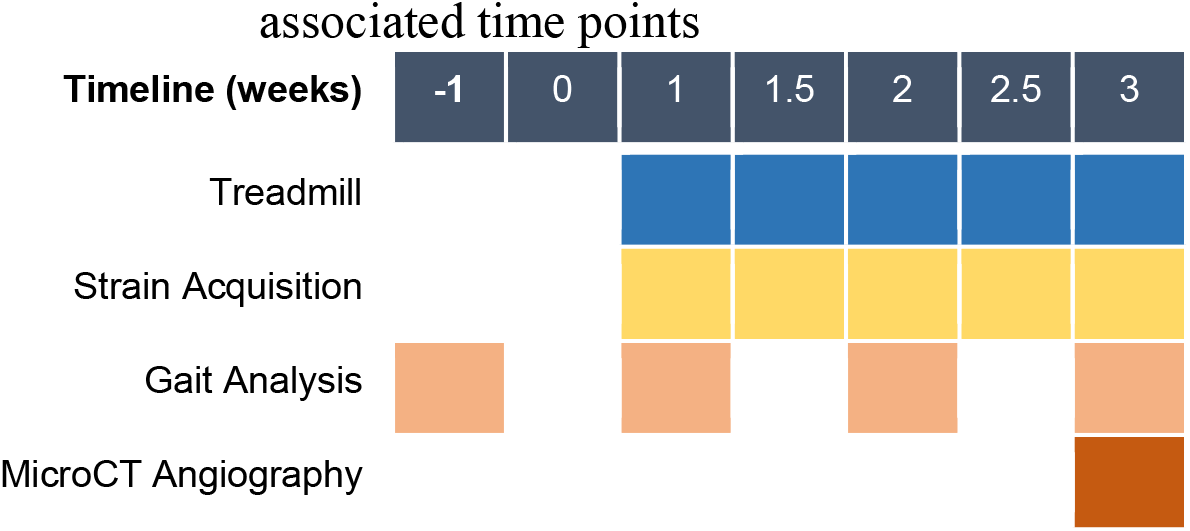
Bone angiography in vivo study timeline. Experimental measurements and associated time points

**Table S3:**
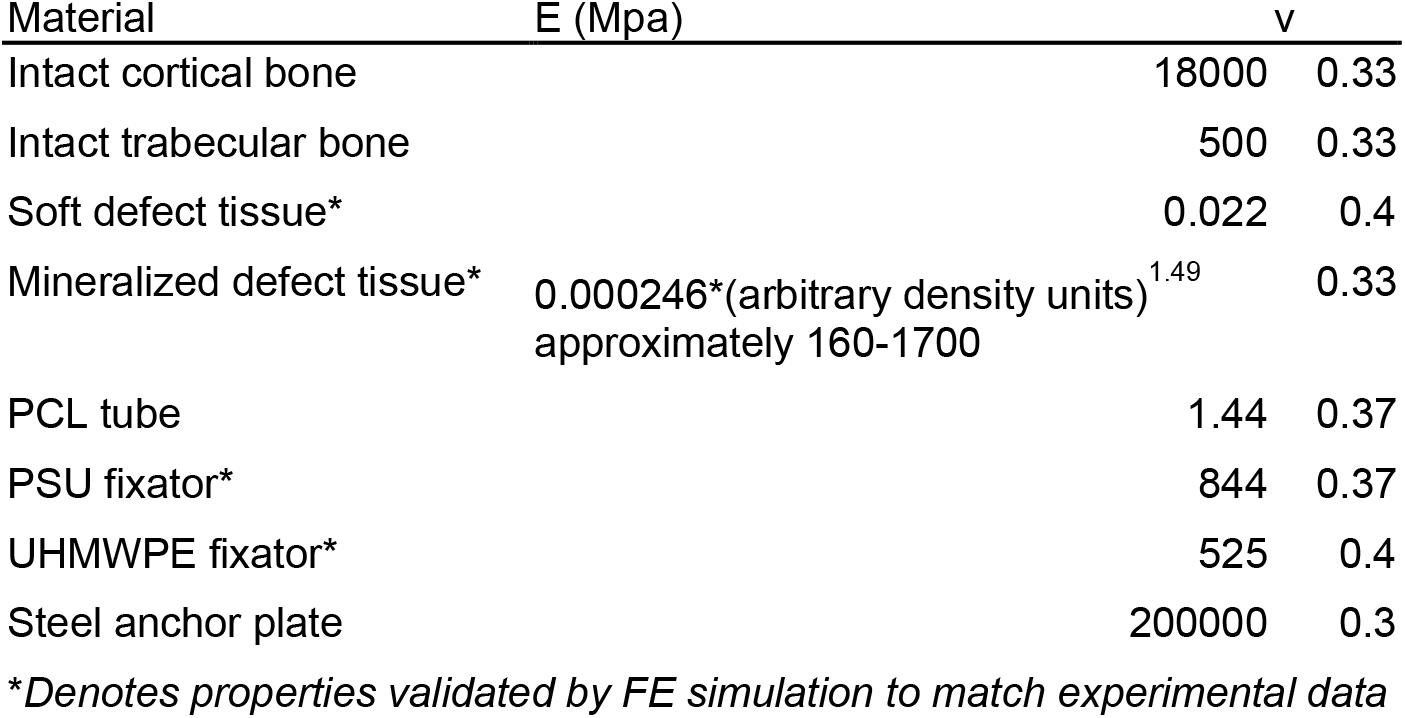
Finite element model mechanical properties.

Movie S1: High-speed X-ray video of real-time strain acquisition during gait 1 month after surgery.

## References and Notes

1. S. W. Lane, D. A. Williams, F. M. Watt, Modulating the stem cell niche for tissue regeneration. Nat. Biotechnol. 32, 795–803 (2014).

2. A. J. Engler, S. Sen, H. L. Sweeney, D. E. Discher, Matrix elasticity directs stem cell lineage specification. Cell. 126, 677–89 (2006).

3. K. H. Vining, D. J. Mooney, Mechanical forces direct stem cell behaviour in development and regeneration. Nat. Rev. Mol. Cell Biol. 18, 728–742 (2017).

4. C. Rot, T. Stern, R. Blecher, B. Friesem, E. Zelzer, A Mechanical Jack-like Mechanism Drives Spontaneous Fracture Healing in Neonatal Mice. Dev. Cell. 31, 159–170 (2014).

5. A. M. McDermott et al., Recapitulating bone development for tissue regeneration through engineered mesenchymal condensations and mechanical cues. Sci. Transl. Med. 11 (2019), doi:10.1101/157362.

6. B. S. Klosterhoff, S. Nagaraja, J. J. Dedania, R. E. Guldberg, N. J. Willett, in Materials and Devices for Bone Disorders, S. Bose, A. Bandyopadhyay, Eds. (Academic Press, ed. 1st, 2017), pp. 197–264.

7. J. D. Boerckel et al., Effects of in vivo mechanical loading on large bone defect regeneration. J. Orthop. Res. 30, 1067–1075 (2012).

8. L. E. Claes et al., Effects of mechanical factors on the fracture healing process. Clin. Orthop. Relat. Res. 355S, S132–47 (1998).

9. R. Zura et al., Epidemiology of Fracture Nonunion in 18 Human Bones. JAMA Surg. 151, e162775 (2016).

10. G. A. Ilizarov, The tension-stress effect on the genesis and growth of tissues. Part I. The influence of stability of fixation and soft-tissue preservation. Clin. Orthop. Relat. Res., 249–81 (1989).

11. A. Pobloth et al., Mechanobiologically optimized 3D titanium-mesh scaffolds enhance bone regeneration in critical segmental defects in sheep. Sci. Transl. Med. 8828 (2018).

12. M. Cilla et al., Machine learning techniques for the optimization of joint replacements: Application to a short-stem hip implant. PLoS One. 12, e0183755 (2017).

13. R. D. Carpenter et al., Effect of porous orthopaedic implant material and structure on load sharing with simulated bone ingrowth: A finite element analysis comparing titanium and PEEK. J. Mech. Behav. Biomed. Mater. 80, 68–76 (2018).

14. E. F. Binder, M. Brown, K. Steger-may, K. E. Yarasheski, K. B. Schechtman, Effects of Extended Outpatient Rehabilitation After Hip Fracture. JAMA. 292, 837–846 (2004).

15. M. Quarta et al., Bioengineered constructs combined with exercise enhance stem cell-mediated treatment of volumetric muscle loss. Nat. Commun. 8, 1–17 (2017).

16. T. A. Rando, F. Ambrosio, Forum Regenerative Rehabilitation : Applied Biophysics Meets Stem Cell Therapeutics. Cell Stem Cell. 22, 306–309 (2018).

17. J. D. Boerckel, B. A. Uhrig, N. J. Willett, N. Huebsch, R. E. Guldberg, Mechanical regulation of vascular growth and tissue regeneration in vivo. Proc. Natl. Acad. Sci. 108, E674–E680 (2011).

18. B. S. Klosterhoff et al., Implantable Sensors for Regenerative Medicine. J. Biomech. Eng. 139, 020806 (2017).

19. P. Nadeau et al., Prolonged energy harvesting for ingestible devices. Nat. Biomed. Eng. 1, 0022 (2017).

20. C. M. Boutry et al., Biodegradable and flexible arterial-pulse sensor for the wireless monitoring of blood flow. Nat. Biomed. Eng. 3, 47–57 (2019).

21. W. T. Abraham et al., Sustained efficacy of pulmonary artery pressure to guide adjustment of chronic heart failure therapy: complete follow-up results from the CHAMPION randomised trial. Lancet. 387, 453–461 (2016).

22. B. S. Klosterhoff et al., Wireless implantable sensor for noninvasive, longitudinal quantification of axial strain across rodent long bone defects. J. Biomech. Eng. 139 (2017), doi:10.1115/1.4037937.

23. J. D. Boerckel et al., Effects of protein dose and delivery system on BMP-mediated bone regeneration. Biomaterials. 32, 5241–51 (2011).

24. C. Schwarz et al., Mechanical Load Modulates the Stimulatory Effect of BMP2 in a Rat Nonunion Model. Tissue Eng. Part A. 19, 247–254 (2013).

25. E. Ozcivici et al., Mechanical signals as anabolic agents in bone. Nat. Rev. Rheumatol. 6, 50–9 (2010).

26. S. M. Perren, Physical and biological aspects of fracture healing with special reference to internal fixation. Clin. Orthop. Relat. Res., 175–96 (1979).

27. G. A. Ilizarov, The tension-stress effect on the genesis and growth of tissues: Part II. The influence of the rate and frequency of distraction. Clin. Orthop. Relat. Res., 263–85 (1989).

28. L. G. Vincent, Y. S. Choi, B. Alonso-Latorre, J. C. del Álamo, A. J. Engler, Mesenchymal stem cell durotaxis depends on substrate stiffness gradient strength. Biotechnol. J. 8, 472–84 (2013).

29. L. Krishnan et al., Effect of mechanical boundary conditions on orientation of angiogenic microvessels. Cardiovasc. Res. 78, 324–32 (2008).

30. G. J. Miller, L. C. Gerstenfeld, E. F. Morgan, Mechanical microenvironments and protein expression associated with formation of different skeletal tissues during bone healing. Biomech. Model. Mechanobiol. (2015), doi:10.1007/s10237-015-0670-4.

31. J. D. Boerckel, K. M. Dupont, Y. M. Kolambkar, A. S. P. Lin, R. E. Guldberg, In Vivo Model for Evaluating the Effects of Mechanical Stimulation on Tissue-Engineered Bone Repair. J. Biomech. Eng. 131, 084502–084502 (2009).

32. C. T. Rubin, L. E. Lanyon, Regulation of bone mass by mechanical strain magnitude. Calcif. Tissue Int. 37, 411–417 (1985).

33. H. Razi et al., Aging Leads to a Dysregulation in Mechanically Driven Bone Formation and Resorption. J. Bone Miner. Res. 30, 1864–1873 (2015).

34. B. W. Hoyt, G. J. Pavey, P. F. Pasquina, B. K. Potter, Rehabilitation of Lower Extremity Trauma : a Review of Principles and Military Perspective on Future Directions. Curr. Trauma Reports. 1, 50–60 (2015).

35. D. L. Miglioretti et al., The use of computed tomography in pediatrics and the associated radiation exposure and estimated cancer risk. JAMA Pediatr. 167, 700–7 (2013).

36. L. E. Claes, J. L. Cunningham, Monitoring the mechanical properties of healing bone. Clin. Orthop. Relat. Res. 467, 1964–71 (2009).

37. M. C. Lin et al., Smart bone plates can monitor fracture healing. Sci. Rep., 1–15 (2019).

38. K. M. Labus et al., Direct Electromagnetic Coupling for Non-Invasive Measurements of Stability in Simulated Fracture Healing. J. Orthop. Res., 1–26 (2019).

39. S. Dalise et al., Biological effects of dosing aerobic exercise and neuromuscular electrical stimulation in rats. Sci. Rep. 7, 1–13 (2017).

40. Y. M. Kolambkar et al., Nanofiber orientation and surface functionalization modulate human mesenchymal stem cell behavior in vitro. Tissue Eng. Part A. 20, 398–409 (2014).

41. Y. M. Kolambkar et al., An alginate-based hybrid system for growth factor delivery in the functional repair of large bone defects. Biomaterials. 32, 65–74 (2011).

42. L. De Visser, R. Van Den Bos, Novel approach to the behavioural characterization of inbred mice : automated home cage observations. Genes, Brain Behav. 5, 458–466 (2006).

43. C. L. Duvall, W. R. Taylor, D. Weiss, R. E. Guldberg, Quantitative microcomputed tomography analysis of collateral vessel development after ischemic injury. Am. J. Physiol. - Hear. Circ. Physiol. 287, H302–H310 (2004).

44. T. Wehner, L. Claes, F. Niemeyer, D. Nolte, U. Simon, Influence of the fixation stability on the healing time--a numerical study of a patient-specific fracture healing process. Clin. Biomech. (Bristol, Avon). 25, 606–12 (2010).

45. D. Ulrich, B. van Rietbergen, A. Laib, P. R̈uegsegger, The ability of three-dimensional structural indices to reflect mechanical aspects of trabecular bone. Bone. 25, 55–60 (1999).

46. T. Wehner et al., Internal forces and moments in the femur of the rat during gait. J. Biomech. 43, 2473–9 (2010).

47. S. A. Maas, B. J. Ellis, G. A. Ateshian, J. A. Weiss, FEBio: Finite Elements for Biomechanics. J. Biomech. Eng. 134, 011005 (2012).

